# Unveiling the effect of phosphorylation on the structural and aggregation properties of the amyloidogenic intrinsically disordered protein DPF3a

**DOI:** 10.1101/2025.08.17.670156

**Authors:** Tanguy Leyder, Julien Mignon, Emma Bongiovanni, Quentin Machiels, Jehan Waeytens, Vincent Raussens, Antonio Monari, Denis Mottet, Catherine Michaux

**Affiliations:** Laboratoire de Chimie Physique des Biomolécules, UCPTS, University of Namur, 5000 Namur, Belgium; Namur Institute of Structured Matter (NISM), University of Namur, 5000 Namur, Belgium; Namur Research Institute for Life Sciences (NARILIS), University of Namur, 5000 Namur, Belgium; Laboratoire de Structure et Fonction des Membranes Biologiques, Université Libre de Bruxelles, 1050 Bruxelles, Belgique; RD3-Pharmacognosy, Bioanalysis and Drug Discovery Unit, Faculty of Pharmacy, Université Libre de Bruxelles, 1050 Bruxelles, Belgique; Université Paris Cité and CNRS, ITODYS, 75006 Paris, France; Gene Expression and Cancer Laboratory, GIGA-Molecular Biology of Diseases, University of Liège, 4000 Liège, Belgium

**Author notes:** Corresponding author at: Laboratoire de Chimie Physique des Biomolécules, UCPTS, Université de Namur, rue de Bruxelles 61, 5000 Namur, Belgique. E-mail address (T. Leyder), (C. Michaux).

**Keywords:** Double PHD fingers 3a (DPF3a), Phosphorylation, Intrinsically disordered protein, Amyloidogenic protein, Spectroscopy, Autofluorescence, Microscopy, Molecular dynamics

## Abstract

The double plant homeodomain fingers 3a (DPF3 isoform a) is a human epigenetic regulator involved in chromatin remodelling, cell division, and ciliogenesis. Most notably, this protein is deregulated in various cancer types and neurodegenerative diseases. In our previous work, the disorder nature of DPF3a, as well as its propensity to aggregate into amyloid fibrils, have been highlighted, making it an amyloidogenic intrinsically disordered protein (IDP). Due to their high chain accessibility, IDPs structure and function are modulated by phosphorylation. It has been reported that phosphorylation of DPF3a at S348 (pS348) by the casein kinase 2 (CK2) is implicated in cardiac hypertrophy. CK2 can also phosphorylate DPF3a at S138 (pS138), which is also located in an intrinsically disordered region (IDR). However, no structural information is available on phosphorylated DPF3a. In the present study, we investigated the effect of phosphorylation on DPF3a structural and aggregation properties. Two single-mutated phosphomimetics (S138E and S348E) were characterised *in vitro* and compared to DPF3a WT, while *in silico* analyses were performed on pS138 and pS348 to assess structural changes at the molecular level. Circular dichroism and fluorescence spectroscopy revealed that both phosphomimetics are hybrid IDPs, with increased turn and antiparallel β-sheet content as well as more buried aromatic residues compared to DPF3a WT, suggesting conformational rearrangements and a more folded N-terminal region. *In silico* characterisation supported these results, showing that phosphorylation of S138 and S348 induce extended conformation, especially the C-terminal extremity, due to electrostatic repulsion, while local folding occurs due to a proximity with arginine and lysine residues. Furthermore, spectroscopic and microscopic analyses unveiled that S138E and S348E exhibit slower fibrillation kinetics compared to DPF3a WT involving distinct aggregation mechanisms.

## 1. Introduction

Posttranslational modifications (PTMs) are essential biochemical processes enhancing the diversity of proteins beyond their genetically encoded sequences and consequently increasing the complexity of biological systems. PTM refers to the covalent attachment of a chemical group to a protein after its synthesis. Many proteins undergo PTMs, and the interplay of these modifications often governs their functions, regulations, interactions, activity, stability, and physiochemical properties. More than 650 PTM types occur in various cellular compartments, with phosphorylation, methylation, acetylation, glycosylation, and ubiquitination being the most common [1–5].

Phosphorylation is the most widespread PTM, affecting approximately a third of the eukaryotic proteome. This reversible kinase-based reaction consists in the addition of a phosphate group from ATP to the side chains of specific amino acids. Phosphorylation primarily targets serine residues (86.4%), followed by threonine residues (11.8%), and then tyrosine residues (1.8%). Conversely, the removal of the phosphate group, known as dephosphorylation, is catalysed by phosphatases [2,6–8]. Protein phosphorylation fulfils several crucial cellular functions, such as growth and development, regulation of cell proliferation, innate, and acquired immunity, as well as differentiation during embryogenesis [9,10]. Such process is involved in aging, which is notably characterised by the abnormal accumulation of senescent cells in tissues and altered neurotransmission [11,12]. Misregulated phosphorylation can alter protein activity, stability, function, and can lead to various severe human pathologies, such as cancers, metabolic disorders, as well as neurodegenerative diseases [2].

It is well-known that phosphorylation particularly occurs in intrinsically disordered proteins (IDPs) and intrinsically disordered regions (IDRs), due to their high flexibility and chain accessibility [13,14]. IDPs are indeed characterised by the lack of a well-defined structure and exist as a dynamic ensemble of interconverting conformers [15,16]. Their singular property makes them more accessible to kinases for sidechain modification. In addition, IDRs contain short linear motifs (SLiMs), which are primary sites for kinase recognition and, consequently, regions where phosphorylation is highly prevalent [17,18]. Phosphorylation regulates IDPs activity either as a molecular switch or a rheostat. It has been widely shown to modulate not only their structural and conformational ensemble, notably by inducing transient secondary structures and disorder-to-order transitions, but also their function and pathogenicity [13,17,19]. For example, phosphorylation of one specific serine residue (S129) of α-synuclein (α-syn) enhances its aggregation into β-sheet-rich amyloid fibrils, thereby influencing its toxicity [20]. The tau protein can undergo hyperphosphorylation preventing its binding to microtubules in the brain, due to the accumulation of negatives charges, thus altering its function and promoting its aggregation into neurofibrillary tangles [21,22].

In the framework of amyloidogenic IDPs, we are interested in the double plant homeodomain (PHD) fingers 3 (DPF3) protein, a human epigenetic regulator belonging to the multiprotein BRM/BRG1-associated factor (BAF) complex. This BAF complex uses the energy of ATP hydrolysis to disrupt DNA-histone interactions, therefore increasing chromatin accessibility [23,24]. A novel BAF-independent function was recently discovered, showing the involvement of DPF3 in mitotic cell division and ciliogenesis [25]. While DPF3 is essential for ciliogenesis through the regulation of axoneme elongation, the protein is dynamically localised in various mitotic structures during mitosis. Knockdown of DPF3 leads to the instability of kinetochore fibres, unstable microtubule-kinetochore attachment, and defect in chromosome alignment, causing mitotic errors, cell death, and genomic instability [25]. From a pathological point of view, DPF3 is not only deregulated in several cancer types [26–30], but may also be implicated in Alzheimer’s disease (AD) and Parkinson’s disease (PD). Indeed, DPF3 contributes to AD development, and elevated expression levels of DPF3 have been detected in neuronal clusters involved in cellular damage associated with PD [31–33]. Furthermore, DPF3 is up regulated in patients suffering from the Tetralogy of Fallot, a cardiac hypertrophy disorder characterised by structural heart defects and right ventricular hypertrophy [34]. In light of its pathophysiological repertoire, DPF3 appears as a new and promising therapeutic target.

In humans, DPF3 exists as two splicing variants, known as DPF3b and DPF3a. The two isoforms are identical from the N-terminal part up to the 292^nd^ residue, including the 2/3 domain, a Krüppel-like zinc finger (ZnF) domain (C_2_H_2_), as well as two IDRs (IDR-1 and IDR-2). In contrast, they differentiate in their C-terminal region: while DPF3b contains two PHD ZnFs (PHD-1 and PHD-2), DPF3a is characterised by a single truncated PHD finger (PHD-1/2), followed by a third IDR (IDR-3) at the C-terminal extremity [23]. Through its PHD tandem, typical of the DPF protein family, DPF3b acts as an epigenetic reader by recognising and binding acetylated and methylated lysine residues on histone tails, allowing the recruitment of the BAF complex and the transcriptional machinery complex for gene transcription [35,36]. Unlike DPF3b, the truncated PHD ZnF domain of DPF3a prevents the binding to modified lysine residues. However, DPF3a contributes to myogenic differentiation through its interaction with the hepatoma-derived growth factor-related protein 2 (HRP2). Specifically, HRP2 binds modified histone and recruits the BAF complex by directly interacting with DPF3a, promoting chromatin accessibility and transcription of myogenic genes [37]. In our previous studies, we have unveiled that both isoforms are hybrid IDPs, with a high disorder content and lacking a hydrophobic core [38], especially in the case of DPF3a, which exhibits even more expanded conformations through its third IDR [24,38,39]. Their propensity to spontaneously aggregate into amyloid fibrils have also been demonstrated characterising both isoforms as amyloidogenic IDPs [38,40].

Through mass spectrometry, DPF3a has been reported to be phosphorylated at S138 and S348 [34]. Interestingly, these residues are respectively located in IDR-1 and IDR-3. S348 is phosphorylated by the casein kinase 2 (CK2) upon hypertrophic stimuli. This allows DPF3a to interact with the transcriptional repressors HEY (HES-related repressor protein), leading to the release of HEY from the DNA and the recruitment of the BAF complex. Hence, the genomic targets, which are foetal genes involved in embryonic cardiac development and are subsequently silenced, are reactivated and transcribed, ultimately leading to cardiac hypertrophy [34]. So, CK2-dependent phosphorylation of DPF3a is essential for its activity. However, no structural information is currently available on phosphorylated DPF3a (pDPF3a) mediated by CK2.

To decipher the impact of the two specific phosphorylations on the structure and the aggregation properties of DPF3a, two single-mutated DPF3a phosphomimetics will first be characterised *in vitro* and their properties compared to wild-type (WT) DPF3a [38]. To this end, S138 and S348 will be mutated into glutamate residues (S138E and S348E), in order to mimic the negative charge and steric hindrance of the phosphate group, and the structural and conformational effects of the mutations will be investigated [3,41]. In parallel, *in silico* analyses will be performed on two monophosphorylated DPF3a (pDPF3a), in which the PTM will be introduced at each of the phosphosites, namely pS138 and pS348, respectively. Importantly, these two phosphorylation sites are located within distinct IDRs of DPF3a, suggesting that they may modulate the protein properties in different ways. Through the use of a combined *in vitro* and *in silico* approach, we aim to analyse the structural and aggregation properties of pDPF3a and compare to those of the WT DPF3a showing the important role of phosphorylation in shaping the conformational landscape of an important epigenetic regulator.

## 2. Materials and methods

### 2.1. Overexpression and purification of DPF3a phosphomimetics

The phosphomimetics S138E and S348E were overexpressed with a GST tag at their N-terminus using a pET-like vector in *E. coli* BL21 (DE3) strains. Transformed bacteria were precultured in 20 g/L lysogeny Lennox broth (LB) containing 0.36 mM ampicillin for 16 h at 37 °C. From 10.0 mL of preculture, strains were cultured in 20 g/L LB Lennox with 0.14 mM ampicillin at 37 °C until the 600 nm-optical density reached 0.5-0.8. Cultures were induced by adding 0.5 mM of isopropyl 𝛽-D-1-thiogalactopyranoside (IPTG) at 37 °C for 4 h. After centrifugation, pellets were recovered and stored at -20 °C. Before purification, pellets were suspended in lysis buffer (phosphate-buffered saline (PBS) pH 7.3, 0.5% Triton X-100, 200 mM KCl, 200 µM phenylmethylsulfonyl fluoride), sonicated in an ice-water bath (6 cycles of 30 s with 30 s pauses). After centrifugation, the supernatant was conserved and 200 µM of PMSF were added. Proteins were purified using an Äkta Purifier fast protein liquid chromatography (FPLC). GST tag proteins were bound to a 5 mL GSTrap pre-packed column using the binding buffer (PBS pH 7.3, 200 mM KCl). The GST tag was cleaved on column at 30 °C for 2 h with the TEV protease in the Tris-buffered saline (TBS) (50 mM Tris-HCl pH 8.0 and 150 mL NaCl). After cleavage, the proteins were eluted in TBS and the column was regenerated with the elution buffer (50 mM Tris-HCl pH 8.0, 20 mM reduced glutathione (GSH)). Protein purity was verified using sodium dodecyl sulphate polyacrylamide gel electrophoresis (SDS-PAGE). For measurements of the spontaneous aggregation properties and amyloid fibrils formation ability, proteins have been incubated at a concentration of 0.2 mg/mL in TBS at ∼20 °C.

### 2.2. UV-visible spectroscopy

Purified proteins were concentrated using a 6-8 kDa cut-off dialysis membrane wrapped in PEG-20000. UV-visible absorption spectra were recorded with a UV-63000PC spectrophotometer (VWR), using a quartz QS cell (Helma) with a 10 nm pathlength. Protein concentration was determined by measuring the absorbance at 214 nm after calculating phosphomimetics molar extinction coefficient at 214 nm using the B. Kuipers and H. Gruppen method (622959 M^-1^.cm^-1^) [42]. After reconcentration, the working protein concentration amounted to 0.2 mg/mL.

### 2.3. Far-UV circular dichroism spectroscopy (far-UV CD)

Far-UV (190-260 nm range) CD spectra were recorded with a MOS-500 spectropolarimeter at 20 °C in TBS, using a 1 mm optical pathlength quartz Suprasil cell (Hellma). Four scans were averaged, buffer baselines were subtracted and corrected spectra were smoothed. The following parameters were used: 15 nm/min scanning rate, 2 nm bandwidth, 0.5 nm data pitch, and 2 s digital integration time. On each spectrum, data are presented as the mean residue ellipticity ([Θ]_MRE_), calculated as follows: [Θ]_MRE_ = (M.θ)/(n-1).(10γ.l), where M is the molecular mass (Da), θ the ellipticity (deg), n the sequence length, γ the protein concentration (mg/mL), and l is the cell pathlength (cm).

### 2.4. Fluorescence spectroscopy

Fluorescence measurements, including intrinsic tryptophan fluorescence (ITF), intrinsic tyrosine fluorescence (ITyrF), and autofluorescence (AF), were performed with an Agilent Cary Eclipse fluorescence spectrophotometer at 20 °C in TBS, using a 10 mm quartz QS cell (Hellma). Emission spectra were recorded from the excitation wavelength of ITF (λ_exc_ = 295 nm), ITyrF (λ_exc_ = 275 nm), and AF (λ_exc_ = 400 nm) up to 600 nm, using the following parameters: 1.0 nm data pitch, 0.1 s averaging time, 10 nm excitation-emission slit width (sw), 600 V photomultiplier tube (PMT) voltage, and 600 nm/min scanning rate. Excitation-emission matrices (EEM) were recorded by varying the excitation wavelength from 200 to 500 nm and the emission wavelength from 200 to 600 nm by a 5.0 nm increment.

### 2.5. Transmission electron microscopy (TEM)

DPF3 aggregates were negatively stained and visualised with a PHILIPS/FEI Tecnai 10 electron microscope at a voltage of 100 kV. Formvar/carbon-coated copper grids were hydrophilised by glow discharge. A droplet of protein was left for 3 min onto the grid, and the excess was eliminated with a piece of blotting paper. The grid was put on a 5 µL droplet of 0.5% (w/v) uranyl acetate for 1 min and air-dried for 5 min before analysis.

### 2.6. Atomic force microscopy-infrared spectroscopy (AFM-IR)

Samples for AFM-IR measurements were diluted at a concentration of ∼0.02 mg/mL. The samples were deposited on freshly cleaved mica. Measurements were conducted in tapping mode with a tap 300 gold coated tips (spring constant 42 N/m and resonance frequency of 300 kHz) on a dry air-purged Bruker Dimension IconIR equipped with a quantum cascade laser (Daylight Solution, San Diego, CA, USA). Throughout the measurements, phase lock loop of both tapping and IR signal were ensured. Images were acquired with 512 lines per sample, at a scan rate of 0.5 Hz. After acquisition of AFM-IR images, spectra were recorded on area of interest on the sample. The IR spectra were corrected by the power spectrum of the QCL. The shown IR spectra are averages of 4 recorded spectra. These spectra were acquired with a resolution of 1 cm^-1^. AFM and IR images were flattened using the NanoScope Analysis software from Bruker. An in-house developed software running on Matlab 7.5.0. was used to treat the recorded IR spectra. Shortly, the baseline of the spectra was corrected and applied a Savitsky-Golay filter to smooth them.

### 2.7. Initial conditions for molecular dynamics (MD) simulations

Given the lack of experimentally resolved structures of the DPF3a isoform (Uniprot ID: Q92784–2), the protein tertiary structure was initially modelled with the MMseqs2-AlphaFold2 approach available on the ColabFold platform and selected according to the first ranked and relaxed model [43,44]. To maintain the folding of the zinc finger C_2_H_2_ motif, the position of the Zn^2+^ cation retrieved from the crystallised C_2_H_2_ domain of DPF2 (PDB ID: 3IUF) [45] was aligned to the DPF3a model, and the protonation state of the corresponding cysteine (CYM) and histidine (HID) amino acids was edited with the PyMOL software [46]. Considering a pH value of 8.0, proper protonation of sidechain atoms was carried out with the ProteinPrepare tool from the PlayMolecule webserver [47]. Introduction of the phosphate group on the serine residues at position 138 and 348 was performed via the Vienna-PTM 2.0 online server [48]. Each phosphorylated DPF3a molecule (pS138 and pS348) was centred in a truncated octahedron water box with a 10 Å buffering water layer from each edge using the *tleap* module implemented in AmberTools. The different systems were subsequently neutralised with the required minimal amount of Na^+^ ions before enforcing a physiological concentration of 150 mM NaCl with respect to the volume of the simulation box.

### 2.8. MD simulation and trajectory analysis

Each phosphorylated and solvated DPF3a system was modelled with the AMBER ff14SB force field [49] and simulated in independent triplicates by all-atom classical molecular dynamics (MD) using the GROMACS 2023.1 suite [50,51]. DPF3a intrinsic disorder was taken into account by applying grid-based energy correction map (CMAP) parameters to its identified

IDRs (residues 90-199, 222-260, and 293-357) [52]. Water molecules parameters were described with the TIP4P-D model [53–55]. For more cost-efficient integration of Newton’s equations of motion, the timestep was increased from 2 to 4 fs by using RATTLE and SHAKE constraints in combination with the Hydrogen Mass Repartition (HMR), for the non-solvent molecules achieved via the *Parmed* package available in AmberTools [56–58]. Additionally, hydrogen bonds were constrained to their proper lengths with the LINCS algorithm [59]. Systems were minimised for a maximum of 50000 steps with the steepest decent algorithm [60], as well as thermalised, equilibrated, and propagated in the isothermal and isobaric (NPT) ensemble. Pressure of 1 bar and temperature of 300 K were maintained constant using the Parrinello-Rahman barostat and a modified Berendsen thermostat (velocity rescaling method), respectively [61,62]. Equations of motion were solved with the leap-frog integrator [63]. Long-range coulombic interactions were determined via the Particle Mesh Ewald (PME) summation [64], and short-range non-bonded interactions were computed using a distance cut-off of 1.0 nm. Production MD simulations were propagated in periodic boundary conditions (PBC) in all three dimensions for an individual duration of 1000 ns. Along the trajectories, atom coordinates and energies were saved every 40 ps.

Regarding trajectory analysis, modules directly implemented in the GROMACS suite and in-house Python scripts were conjunctively used. Time-dependent variables were computed and averaged from the resulting triplicates for each phosphorylated protein. Whereas root-mean square deviation (RMSD) was determined from the backbone atoms of the minimised and equilibrated structure, root-mean square fluctuation (RMSF) gave access to backbone flexibility after alignment of the time-averaged structure taken as reference. Conformational landscape of the protein was represented by two-dimensional bivariate Kernel density graphs of the radius of gyration (R_g_) against the end-to-end distance (d_ee_). Considering the protein as a whole, solvent accessible surface area (SASA), number of contacts within 5 Å, and intramolecular H bonds within a 3.5 Å donor-acceptor distance and 30° angle cut-off were calculated. A total of 3000 frames was clustered for each phosphorylated system using a RMSD cut-off of 5 Å between the nearest structural neighbours, and the central structure of the most populated cluster amongst the replicates was extracted to map the minimum distances for every pair of residues. Secondary structure classes and analysis were defined with the STRIDE algorithm in combination with the Timeline VMD plugin [65]. Seven classes of secondary structures were assessed: turns (predominantly β type, *i-i+3* turn), extended β-sheet, isolated β-bridge (usually *i-i+>6*), α-helix (*i-i+5* helix), 3_10_-helix (*i-i+3* helix), π-helix (distorted or bulged *i-i+5* helix), and random coil. Visualisation of the trajectories and rendering of selected snapshots were achieved with the VMD software [66].

## 3. Results and discussion

### 3.1. Structural properties of DPF3a phosphomimetics

In order to investigate the effect of phosphorylation on the structure of DPF3a, various biospectroscopies were applied on the two designed phosphomimetics (S138E and S348E), and the results have been compared to that of DPF3a WT previously published [38]. Firstly, far-UV circular dichroism (CD) spectroscopy was performed to evaluate the presence of secondary structure elements in DPF3a proteins. Similarly to DPF3a WT, the two phosphomimetics are hybrid IDPs (Fig. 1). Indeed, while the maximum around 200 nm is associated to ordered secondary structures, the first minimum near 206 nm is indicative of disorder. As a matter of fact, a fully disordered protein typically displays a minimum at shorter wavelengths, around 200 nm, with no maximum in the 190-200 nm range [67,68]. While the CD spectrum of DPF3a WT presents a broad shoulder around 225 nm, characteristic of the presence of a small content of antiparallel β-sheets, spectra of the phosphomimetics reveal a more pronounced second minimum appearing at this wavelength, suggesting that S138E and S348E contain a higher proportion of antiparallel β-sheets. Such disorder-to-order transition upon phosphorylation has already been observed for other IDPs [3,14]. Moreover, the maximum observed between 210 and 220 nm indicates an enrichment in turns in both phosphomimetics compared to DPF3a WT [69–71]. Nevertheless, the two phosphomimetics contain less α-helix with respect to DPF3a WT. The minimum shifted at 207 nm and the broader shoulder between 220 and 225 nm in the DPF3a WT spectrum could correspond to α-helices, typically characterised by two distinct minima at 208 and 222 nm [72].

**Fig. 1.**
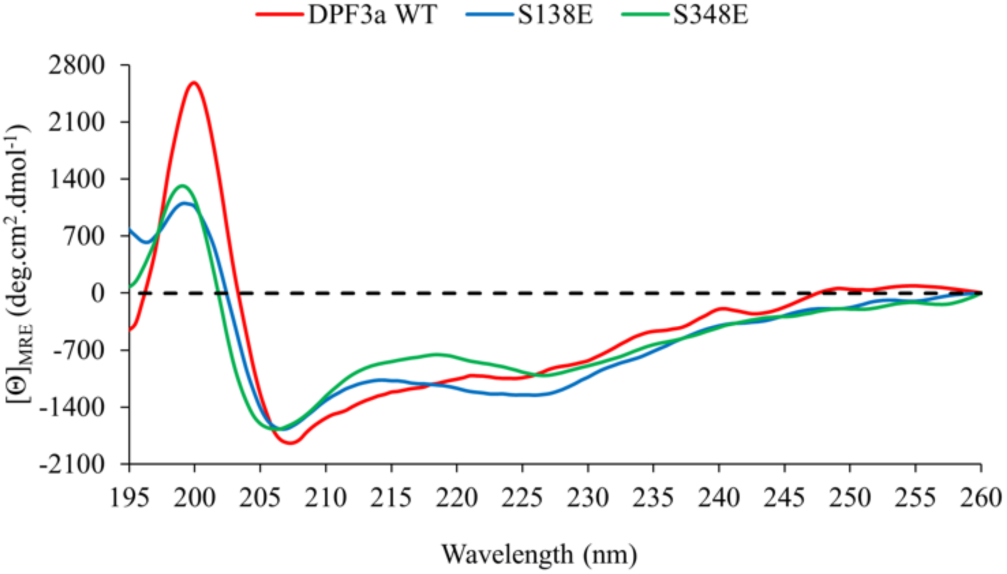
Far-UV CD spectra of DPF3a WT (red), S138E (blue) and S348E (green) in TBS at ∼ 20 °C.

Secondly, local conformations of the phosphomimetics and their folding state were investigated by fluorescence spectroscopy. Indeed, the exposure of solvatochromic tryptophan (Trp) and tyrosine (Tyr) residues has been probed by intrinsic tryptophan fluorescence (ITF) and intrinsic tyrosine fluorescence (ITyrF). The emission band position of Trp residues is influenced by its direct environment, giving rise to characteristic emission wavelengths varying from 308 to 355 nm after excitation at 295 nm. Specifically, the emission shifts to red or the blue in polar or hydrophobic environments, respectively. More precisely, fully solvent-exposed, partially exposed, and buried Trp residues usually emit at ∼350 nm, around 330-340 nm, and between 308 and 315 nm, respectively [73]. The ITF spectrum of DPF3a WT displays an emission band at ∼340 nm with a slight shoulder at higher wavelengths relative to Trp residues partially exposed to the solvent and/or polar amino acids (Fig. 2A). DPF3a contains two Trp residues (Trp^56^ and Trp^79^) in its primary structure, both located within the 2/3 domain which is predicted to be predominantly ordered while retaining some flexibility [38]. Comparatively, the Trp emission bands of S138E and S348E are found at ∼338 and ∼330 nm, respectively. This hypochromic effect is explained by less solvent-exposed Trp residues, which is exacerbated in the case of S348E. Both spectra do not present a shoulder at higher wavelengths confirming that Trp residues are more buried. Therefore, it is expected that phosphorylation, especially at position 348, induces either locally a higher degree of folding in the 2/3 domain or the overall burying of Trp residues into more compact conformers. Interestingly, the ITF spectrum of DPF3b, which is more ordered than DPF3a, also presents an emission band at ∼335 nm [38].

**Fig. 2.**
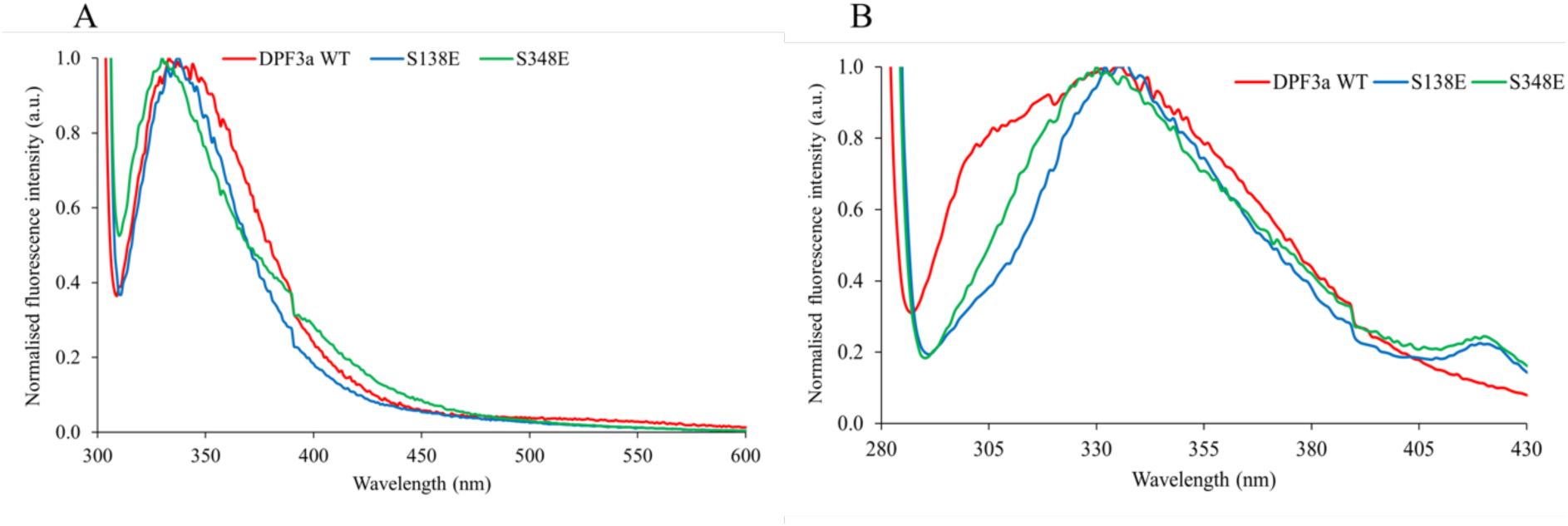
Intrinsic fluorescence of DPF3a WT, S138E and S348E in TBS at ∼ 20 °C. (A) Normalised ITF spectra (λ_exc_ = 295 nm, sw = 10 nm) of DPF3a WT (red), S138E (blue) and S348E (green). (B) Normalised ITyrF spectra (λ_exc_ = 275 nm, sw = 10 nm) of DPF3a WT (red), S138E (blue) and S348E (green).

Complementarily, ITyrF allows the investigation of the local environment of Tyr residues, although they are less sensitive to environmental changes. Upon excitation at 275 nm, their emission band typically appears between 300 and 310 nm. However, Trp-Tyr fluorescence resonance energy transfer (FRET) usually hides ITyrF signature due to Trp and Tyr spectral overlapping, depending on their sequence position, and spatial proximity [74]. DPF3a contains 10 Tyr residues, distributed along its sequence as follows: 5 within the 2/3 domain and close to the two Trp residues (Tyr^17^, Tyr^27^, Tyr^54^, Tyr^72^, and Tyr^74^), 4 in the C_2_H_2_ zinc finger (Tyr^198^, Tyr^207^, Tyr^215^, and Tyr^217^), and 1 found in the PHD-1/2 domain (Tyr^261^). DPF3a WT ITyrF emission spectrum shows a principal emission band centred at ∼338 nm, originating from Trp-Tyr FRET, along with a shoulder around 305 nm, associated with freely emitting Tyr residues exposed to the solvent (Fig. 2B). While the Trp-Tyr FRET emission band appears at ∼337 nm and ∼332 nm for S138E and S348E, respectively, in good agreement with ITF results, the shoulder around 305 nm is less pronounced for both phosphomimetics compared to DPF3a WT and is nearly absent. This suggests that, comparatively to the Trp residues, the Tyr residues are globally less exposed to the solvent compared to those in DPF3a WT and that Trp-Tyr FRET is enhanced for both phosphomimetics, which is consistent with increased residue bury upon DPF3a phosphorylation.

Regarding these results, it indicates that the phosphomimetics induce local conformational changes, with more ordered regions where Trp and Tyr residues appear more buried compared to DPF3a WT. A similar phenomenon has already been observed for the tau protein where phosphorylation of specific residues can induce local structural rearrangements and the formation of intramolecular interactions, contributing to regional ordering and potentially affecting its interaction with microtubules [75].

### 3.2. Conformational dynamics of monophosphorylated DPF3a molecules

To gain insight into the influence of each phosphorylated variant on the global and local conformational properties of DPF3a, all-atom classical molecular dynamics (MD) simulations were carried out by introducing a phosphate group on serine residues at position 138 (pS138) and 348 (pS348). The MD simulations involving phosphorylation are hereafter analysed and compared to the trajectories of the WT protein which have been obtained using the same force field, water model, and simulation parameters as previously published in [39].

First, the root-mean square deviation (RMSD) evolution over time shows that, irrespectively to the phosphorylation state, all three proteins exhibit substantial deviation from the starting modelled structure within the first 100 ns, the amplitude of which is typically observed for disordered and dynamical polypeptide chains (Fig. 3A). Nevertheless, the presence of a phosphate moiety seemingly leads to more structural heterogeneity, as evidenced by a more fluctuating plateau. Larger variability between replicates is also visible for pS138 in the second half of the simulation. Notably, the RMSD of pS138 and pS348 generally remain smaller throughout the simulation, indicating less deviations from the initial structure and presumably more stable structures compared to DPF3a WT.

**Fig. 3.**
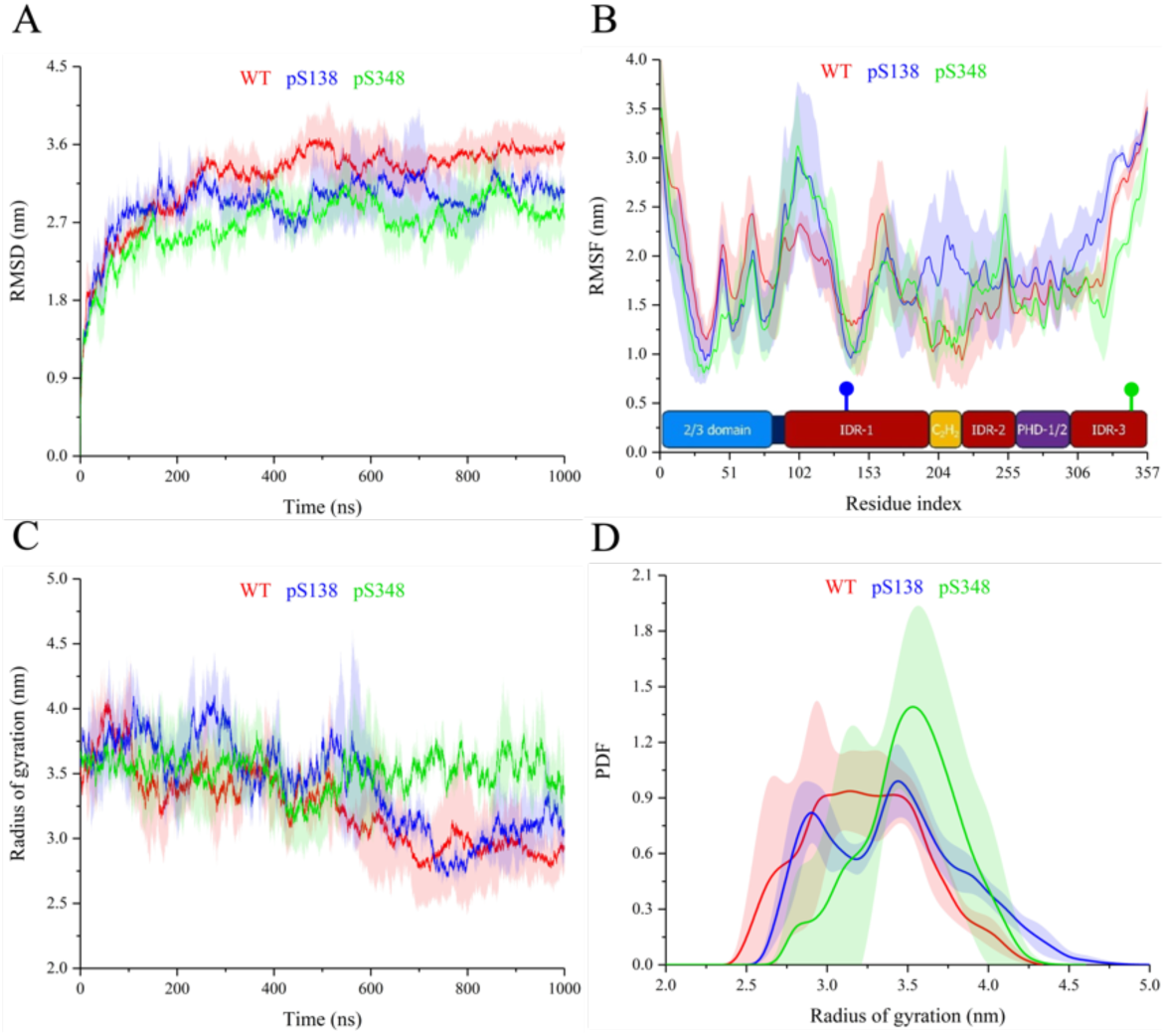
Time-evolution of (A) RMSD and (B) backbone RMSF of full-length WT (red), pS138 (blue), and pS348 (green) DPF3a simulated for 1 µs at pH 8.0 and 150 mM NaCl. For each system, curves correspond to the average of triplicates with the standard deviation represented as a trace (shaded area) in the corresponding colour. At the bottom of the RMSF graph, the sequence organisation is displayed with respect to its constitutive domains: the N-terminal 2/3 domain (blue), intrinsically disordered regions (dark red), Krüppel-like C_2_H_2_ zinc finger (yellow), and truncated PHD-1/2 (purple). Phosphosite S138 and S348 localisation is indicated by a blue and a green stick, respectively. (C) Time-evolution and (D) probability density function (PDF) distribution over the trajectories of the radius of gyration of full-length WT (red), pS138 (blue), and pS348 (green) DPF3a simulated for 1 µs at pH 8.0 and 150 mM NaCl.

Regarding changes in time-averaged root-mean square fluctuation (RMSF) at the backbone level, the overall profile remains similar (Fig. 3B). Nevertheless, scattered variations along the sequence uncover local phosphorylation-dependent effects. While the N-terminal 2/3 domain slightly becomes more rigid upon phosphorylation, consistent with the two Trp residues being more buried for S138E, to a greater extent, for S348E, the flexibility of the first half of IDR-1 is conversely increased. For pS138 only, the C_2_H_2_ ZnF significantly increases its flexibility with nonetheless important variability between replicates. Indeed, pS138 retains this enhanced flexibility up to the C-terminal extremity. Interestingly, the pS138 chain flexibility decreases at the corresponding S138 site and neighbouring amino acids in IDR-1, which is surprisingly also observed for pS348. In comparison, a global stiffening of IDR-3 is induced in pS348, where the phosphosite is located, whilst the region becomes more flexible upon phosphorylation at S138.

Examination of the chain compaction state from the point of view of the radius of gyration (R_g_) reveals that pS138 follows a similar trend to that of WT DPF3a, with R_g_ values steadily decreasing and seemingly plateauing towards the end of the simulation (Fig. 3C). However, the chain sporadically expands along the trajectory, e.g. between 200 and 350 ns, hence populating a second and well-defined R_g_ maximum at around 3.4 nm, while the first one is situated at ∼ 2.8 nm (Fig. 3D). While a more pronounced decrease is observed around 450 ns, phosphorylation of S348 maintains DPF3a in a more swollen conformation throughout the trajectories, which are less subjected to overtime fluctuations – a behaviour that markedly differs from that of WT DPF3a – and results in a narrower R_g_ distribution centred at 3.5 nm. Overall, both phosphorylated variants also populate high R_g_ values, i.e. beyond the 4.0 nm range, compared to WT DPF3a, which has a broader R_g_ distribution towards low R_g_ values. This indicates that the phosphomimetics tend to adopt more extended conformations.

The conformational landscape of the three different forms of DPF3a can be examined by plotting the correlation between the R_g_ and the end-to-end distance (d_ee_) into two-dimensional Kernel density maps. The R_g_-d_ee_ spaces is quite similar for WT DPF3a (Fig. 4A) and pS138 (Fig. 4B), as their Rg values remain relatively close during the simulation. Although both proteins can adopt relatively swollen conformations, certain populations of pS138 exhibit higher R_g_ and d_ee_, compared to DPF3a WT, which is in line with the previous results, particularly the increased RMSF of the C-terminal region and sporadic chain expansion. Phosphorylation of S348 leads to a narrower range of populated conformations, characterised by similar d_ee_ but high R_g_ values, as previously observed (Fig. 4C). Such restriction of the conformational subspace substantiates that the introduction of a phosphate moiety at position 348 induces a consistent global swelling of DPF3a structure. The different populations of the three proteins exhibit d_ee_ values within the same range (4-7 nm), suggesting no significant spatial proximity between the C- and N-terminal extremities, irrespective to the phosphorylation state.

**Fig. 4.**
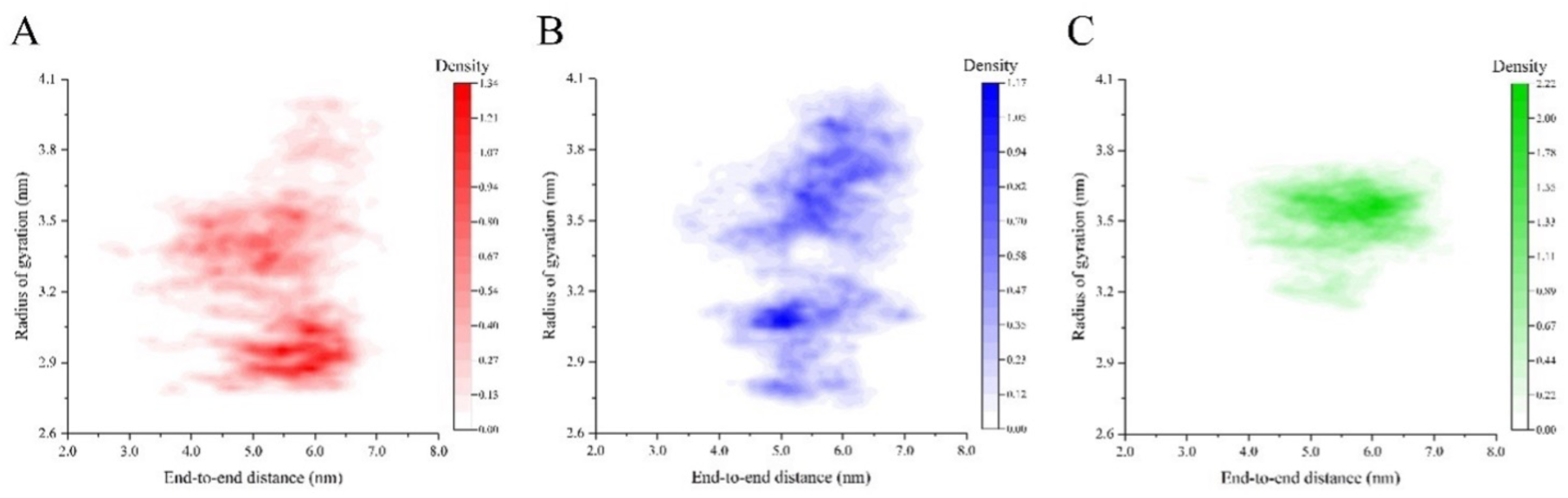
Conformational landscape of full-length (A) WT (red), (B) pS138 (blue), and (C) pS348 (green) DPF3a simulated for 1 µs at pH 8.0 and 150 mM NaCl. For each system, radius of gyration and end-to-end distance pairs were averaged between the triplicates and represented on a two-dimensional Kernel density map in the corresponding colour.

Consistently, DPF3a WT solvent accessible surface area (SASA) steadily decreases over time (Fig. S1A), resulting in a broad SASA distribution centred at 333 nm^2^ (Fig. S1B), with high reproducibility between replicates. pS138 follows the same trend, with SASA values consistently remaining slightly above those of DPF3a WT, except for the first 200 ns, confirming that the primary population is centred around 338 nm^2^. However, greater variability is observed between replicates. In contrast, the SASA profile of pS348 remains relatively constant over time, except for a rapid decrease during the first 100 ns. Consequently, SASA values are lower than those of WT DPF3a and pS138 up to approximately 500 ns, before becoming higher towards the end of the simulation. This results in a narrower population with a maximum around 336 nm^2^. Surprisingly, pS348 exhibits a higher averaged R_g_ compared to pS138, but a smaller averaged SASA. Such divergence suggests that phosphorylation of S348 induces conformational changes leading to a more extended structure (reflected by an increased R_g_), which nonetheless involves local burying of specific regions, thereby contributing to a reduced SASA. As such, the decreased SASA might not reflect a global compaction, but rather a localised rearrangement of the protein.

The phosphorylation of DPF3a, introducing two negative charges, is assumed to facilitate the interactions between the phosphoserine and positively charged residues, most likely influencing the protein conformational state. The radial distribution function g(r) profile between (phospho)serine (Oε)Oγ and arginine Nη atoms (Fig. 5A), as well as lysine Nζ atoms (Fig. 5B), reveals a major population at approximately 0.3 nm for pS138 and pS348, which is indicative of interactions between the oxygen atoms of the phosphate group and the positively charged Arg and Lys residues. Such population is absent or significantly smaller for non-phosphorylated serine residues of WT DPF3a, highlighting the role of phosphorylation in promoting these specific electrostatic interactions. The proximity of certain positively charged residues with respect to the phosphoserines could locally stabilise some regions, thus contributing to a decrease in SASA through the promotion of local compaction.

**Fig. 5.**
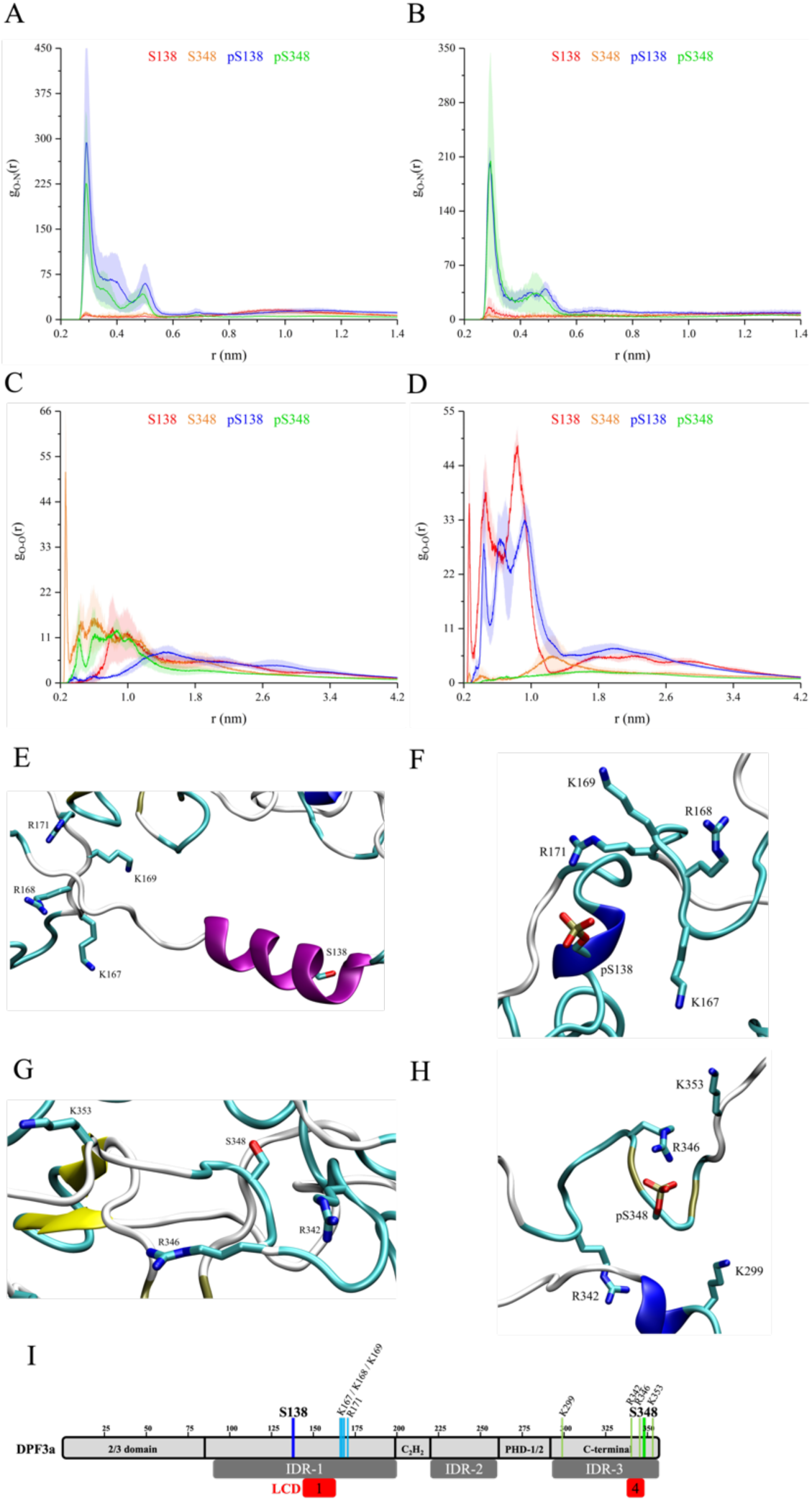
Radial distribution function g(r) between (phospho)serine (Oε)Oγ and (A) arginine Nη atoms, (B) lysine Nζ, (C) aspartate Oδ atoms, and (D) glutamate Oε atoms for WT S138 (red), WT S348 (orange), pS138 (blue), and pS348 (green). For each system, curves and scattered dots correspond to the average of triplicates with the standard deviation represented as a trace (shaded area) in the corresponding colour. Zoom on (E) S138, (F) pS138, (G) S348, and (H) pS348, highlighting neighbouring arginine and lysine residues. For each system, the inset corresponds to the central protein structure of the most populated cluster amongst the triplicates, which is shown in cartoon representation with the different secondary structure elements coloured according to STRIDE assignment: turn (turquoise), extended β-sheet (yellow), isolated β-bridge (tan), α-helix (pink), 3_10_-helix (blue), π-helix (red), and random coil (light grey). (I) Distribution and location of S138 (dark blue) and nearby positively charged residues upon phosphorylation (light blue), S348 (dark green) and nearby positively charged residues upon phosphorylation (light green), the predicted IDRs (dark grey), LCD-1 and LCD-4 (red) along DPF3a sequence.

The introduction of two negative charges may also influence electrostatic repulsion with nearby negatively charged residues. The radial distribution function g(r) between (phospho)serine (Oε)Oγ and aspartate Oδ atoms (Fig. 5C), as well as glutamate Oε atoms (Fig. 5D) provides valuable insight into the impact of such repulsion to the conformational changes observed upon DPF3a phosphorylation, as the protein contains 23 Asp and 41 Glu residues. For both phosphorylated systems, the negatively charged residues tend to move away from the serine upon phosphorylation. This is particularly pronounced for Asp and Glu residues around phosphorylated S138 and S348, respectively. Interestingly, for S138, a significant short distance population with a proximity of Glu residues is still observed even after phosphorylation. This is due to the localisation of S138 near a low complexity domain (LCD) enriched in Glu residues (LCD-1; residues 145 to 164) (Fig. 5I) [38]. However, the population around 0.25 nm for the unmodified S138 disappears after phosphorylation, indicating a loss of short-range interactions. This confirms that phosphorylation induces electrostatic repulsion, potentially likely promoting an extended protein conformation and thereby contributing to the increased R_g_ observed for phosphorylated DPF3a.

In order to visualise and better understand the interactions involving the (phospho)serine and charged residues, we examined a zoomed-in snapshots of the (phospho)serine residues in the central protein structures of the most populated cluster amongst the triplicates. In the case of pS138 (Fig. 5F), a pronounced reorientation of the positively charged stretch, notably encompassing K167 and R171, is observed, bringing it into close vicinity with the phosphoserine compared to the WT structure (Fig. 5E). Moreover, the α-helix comprising S138 undergoes a local structural rearrangement upon phosphorylation, adopting a combination of 3_10_-helix and turn conformations. For S348, even in the absence of phosphorylation (Fig. 5G), the residue is located near at the edge of a LCD enriched in Arg residues (LCD-4; residues 337 to 348) (Fig. 5I) [38], making it directly neighbouring a positively charged environment. Upon phosphorylation, such proximity is spatially enhanced, with a clear repositioning of surrounding Arg residues towards the phosphoserine. R346 forms a salt bridge with phosphorylated S348, while K299, which is not in proximity in the unphosphorylated DPF3a, is evidently reoriented towards pS348 (Fig. 5H), further contributing to the local rearrangement around the phosphoserine.

The phosphorylation of S138 and S348 may also change the occurrence of H bonds within the protein structure. For WT DPF3a, the number of H bonds increases over time as the structure gradually compacts (Fig. 6A), reaching an average of 164 H bonds across the trajectories (Fig. 6B). The same trend is observed for pS138, albeit with a lower number of H bonds count, averaging around 155. In contrast, apart from the first 100 ns, the H bond count in pS348 progressively decreases throughout the simulation, leading to a broader distribution centred around 162. The proximity between phosphoserine and positively charged residues can facilitate the H bonds formation around the phosphoserine. Nonetheless, the overall reduced number of H bonds aligns with the higher R_g_ values observed for the phosphorylated forms of DPF3a, indicating increased conformational flexibility and reduced compactness. This could notably be explained by the electrostatic repulsion between the phosphate group and nearby Asp and Glu amino acids.

**Fig. 6.**
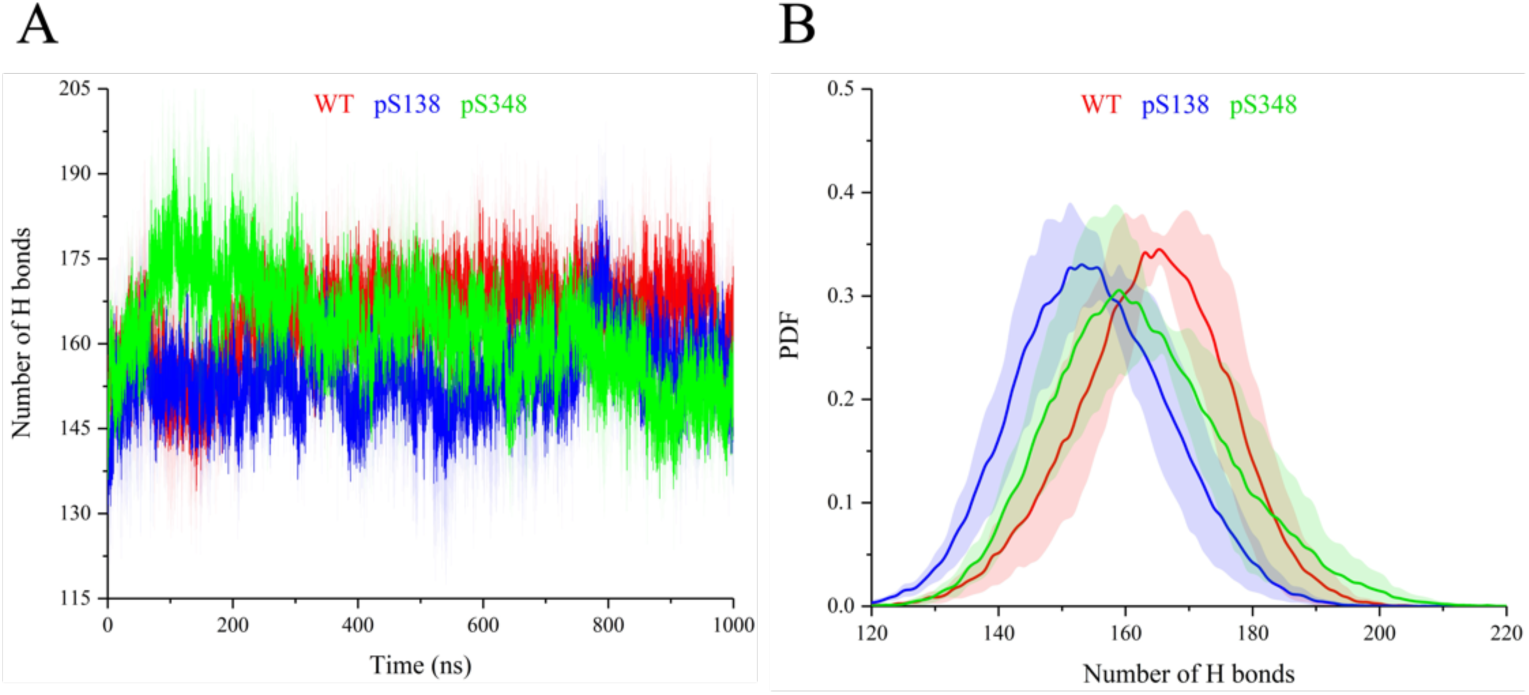
Intramolecular protein H bond (A) evolution over time and (B) probability density function (PDF) distribution over the trajectories of full-length WT (red), pS138 (blue), and pS348 (green) DPF3a simulated for 1 µs at pH 8.0 and 150 mM NaCl. For each system, curves correspond to the average of triplicates with the standard deviation represented as a trace (shaded area) in the corresponding colour.

To further analyse the conformation of the proteins, minimum distance contact maps between residue pairs were compared between the two phosphorylated systems with respect to the reference WT DPF3a as a reference. As for the described snapshots, these maps were determined with the central structure corresponding to the most populated cluster amongst the triplicated trajectories. The structures of pS138 and pS348 are more extended than that of DFP3a WT, which explains their higher R_g_ values, particularly for pS348. While DPF3a WT is in a compact state with every domain interacting with the others, phosphorylation of S138 induces a chain extension, leading to reduced interactions between the different regions. This extension is particularly evident in the C-terminal extremity, which is clearly visible in the represented structure (Fig. 7A). This is highlighted by a loss of interactions and an increased distance between the IDRs, as well as between the IDRs and the other domains, particularly the IDR-3 which become more isolated. The same results are observed for pS348, with the IDRs being separated and the C-terminal region being even more extended, despite a local folding around the phosphoserine (Fig. 7B). However, the C_2_H_2_ ZnF is less isolated and maintains some contacts with the IDR-1, which is in line with the RMSF profile. Additionally, for both phosphorylated proteins, new contact regions (indicated by the black arrows) emerge around the phosphoserines, corresponding to Arg-Lys clusters surrounding each respective phosphosite. Such positively charged environments enable electrostatic interactions with the phosphate group, promoting local folding and thereby contributing to a SASA reduction.

**Fig. 7.**
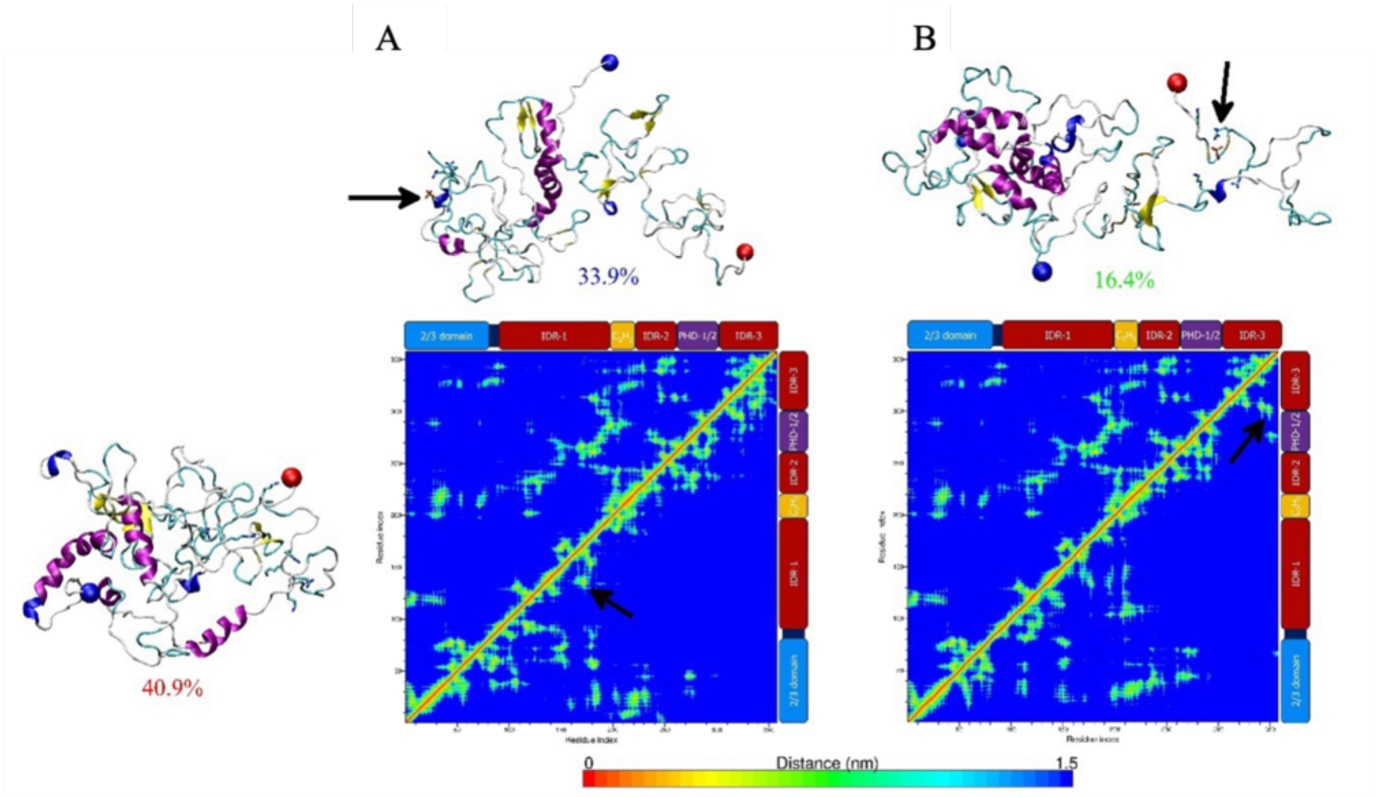
Conformational clustering analysis and residue pairs contact mapping of full-length WT (bottom left), (A) pS138, and (B) pS348 DPF3a simulated for 1 µs at pH 8.0 and 150 mM NaCl. For each system, the central protein structure of the most populated cluster amongst the triplicates is shown in cartoon representation with the N- and C-termini respectively pinpointed as blue and red spheres, and the different secondary structure elements coloured according to STRIDE assignment: turn (turquoise), extended β-sheet (yellow), isolated β-bridge (tan), α-helix (pink), 3_10_-helix (blue), π-helix (red), and random coil (light grey). The coordinated Zn^2+^ cation is represented as a grey bead. Percentages indicate the population size of the first cluster over the selected trajectory. Minimum distance contact maps between residue pairs are compared between the two phosphorylated systems (bottom half on each map) with respect to the reference WT protein (upper half on each map). On the sides of each contact map, the sequence organisation of DPF3a is displayed according to its constitutive domains: the N-terminal 2/3 domain (blue), intrinsically disordered regions (dark red), the Krüppel-like C2H2 zinc finger (yellow), and truncated PHD-1/2 (purple). Black arrows pinpoint the localisation of phosphoserine amino acids and their respective Arg-Lys cluster on the representative snapshots and contact maps.

These phosphorylation-induced structural changes could affect DPF3a function and its interaction with binding partners. Indeed, previous studies have shown that the specific interaction between DPF3a and the basic helix-loop-helix transcription factor HEY1 is facilitated by the DPF3a-specific PHD-1/2 domain and/or its C-terminal region. Notably, phosphorylation of S348 has been identified as critical for the DPF3a-HEY1 interaction [34]. Given that S348 phosphorylation modulates the structure of the C-terminal extremity, it is plausible that this PTM may influence and regulate the interaction with HEY1.

The evolution of secondary structure content allows comparing the conformation and the structure of pS138 (Fig. 8B) and pS348 (Fig. 8C) with the one WT DPF3a (Fig. 8A) [39]. Similarly to WT DPF3a, no noticeable modification of extended β-sheet, isolated β-bridge, 3_10_-helix, and π-helix is observed for pS138 and pS348 during the simulation, with their abundance remaining almost always below 5%. However, both phosphorylated forms of DPF3a WT exhibit a decrease in the α-helix content, whose abundance remain lower than that of WT DPF3a, with values of 11.7% for pS138 and 16.6% for pS348, compared to 16.7% for WT DPF3a. This reduction may also account for the loss of H-bonds observed during the simulation. Both pS138 and pS348 exhibit a similar proportion of coil (approximately 40%), but an enrichment in turn content compared to WT DPF3a, with values of 37.8% for pS138 and 35.6% for pS348, versus 33.8% for the WT. All these analyses are consistent with the experimental CD spectra. In addition, similarly to WT DPF3a, pS138 shows an increase in coil content during the simulation, compensating for the loss of α-helix. A similar trend is observed for pS348, but with an additional increase in turn content.

**Fig. 8.**
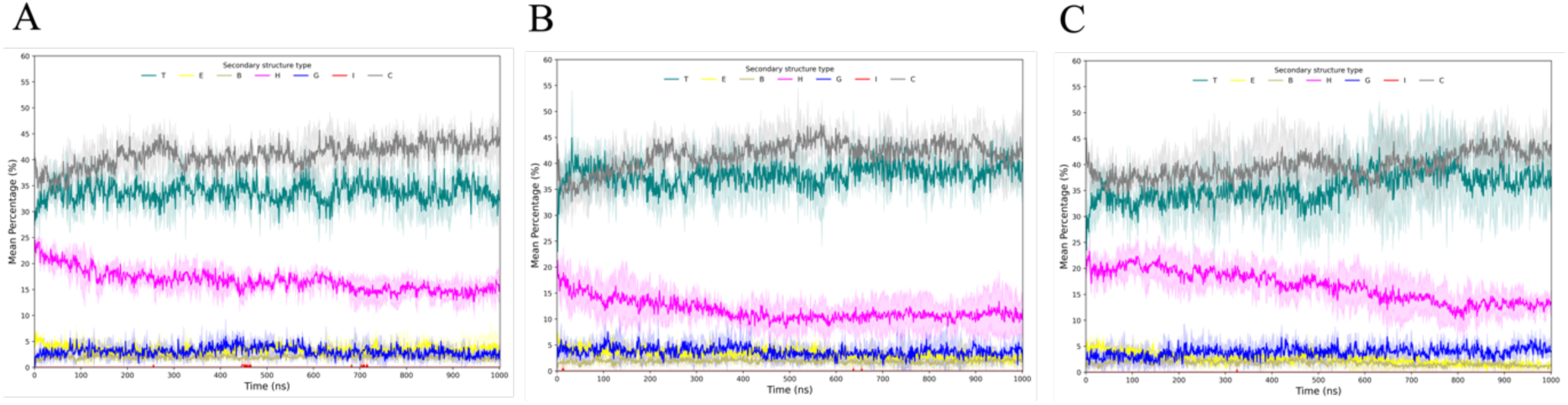
Time-evolution of secondary structure content of full-length (A) WT (B) pS138 and (C) pS348 DPF3a. The secondary structures are defined into seven classes after STRIDE assignment: turn (T, turquoise), extended β-sheet (E, yellow), isolated β-bridge (B, tan), α-helix (H, pink), 3_10_-helix (G, blue), π-helix (I, red), and random coil (C, grey). For each secondary structure, curves correspond to the average of triplicates with the standard deviation represented as a trace (shaded area) in the class-associated colour.

Taken together, these analyses reveal that phosphorylation at S138 and S348 promotes more extended conformations, particularly at the C-terminal extremity, likely driven by electrostatic repulsion. At the same time, local folding is observed, presumably due to the proximity of positively charged residues.

### 3.3. Effect of phosphorylation on the aggregation properties

#### 3.3.1. Spectroscopic analysis of DPF3a phosphomimetics aggregation properties

DPF3a being an amyloidogenic IDP [38], the impact of phosphorylation on its spontaneous aggregation properties and amyloid fibrillation propensity has been investigated. CD is a useful tool to follow the secondary structure modifications during the aggregation process, as well as to establish an aggregation mechanism [76]. Similarly to WT DPF3a (Fig. 9A), the CD spectra of S138E (Fig. 9B) and S348E (Fig. 9C) exhibit signal changes over the course of 96 h. WT DPF3a undergoes conformational rearrangement within 24 h, seemingly resulting in the formation of α-helix-enriched intermediates visualised by the minima at 207 and 220 nm. The two phosphomimetic proteins present similar signature over time. After 24 h of incubation, while the maximum at 200 nm presents no significant shift, a decrease of the intensity of the bands around 206 and 225 nm, corresponding respectively to enrichment in disorder and antiparallel β-sheets, is observed. S138E and S348E present a distinct aggregation mechanism, compared to WT DPF3a, involving the formation of turn intermediates characterised by a maximum at ∼215 nm and the decrease in the intensity of the bands at 206 and 225 nm [69–71]. The subsequent 24 h are characterised by additional modifications akin to those observed for WT DPF3a, i.e. the loss of the disorder-associated band at 206 nm, as well as the clear appearance of a broad negative band at ∼225 nm, indicative of an increase in antiparallel β-sheets. This could be associated with the formation of amyloid fibrils, as observed for WT DPF3a and WT DPF3b, and other amyloidogenic IDPs [77–79]. No further conformational changes occur up to 72 h of incubation, at the exception of a steady enrichment in antiparallel β-sheets, which is even more pronounced for S348E.

**Fig. 9.**
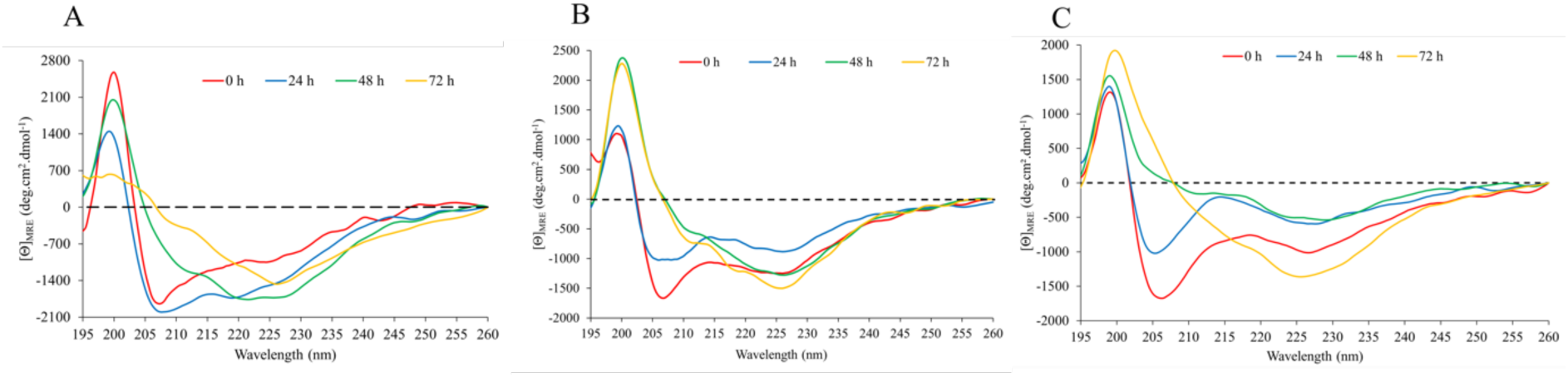
Far-UV CD spectra of (A) DPF3a WT, (B) S138E and (C) S348E after 0 h (red), 24 h (blue), 48 h (green), and 72 h (yellow) of incubation in TBS at ∼ 20 °C.

Regarding ITF spectrum of S138E (Fig. 10B), the emission band initially present at 338 nm progressively shifts towards shorter wavelengths, reaching 333 nm after 72 h of incubation. This is indicative of more buried Trp residues in a hydrophobic core and consistent with the aggregation of the protein. The same tendency is observed for WT DPF3a (Fig. 10A), where the emission band is blue-shifted from 340 nm to 332 nm after 72 h, accompanied by the loss of the shoulder at higher wavelengths. Nonetheless, on WT DPF3a spectra, the hypsochromic shift is accompanied by an hyperchromic effect throughout the aggregation, related to Trp residues found in a more non-polar environment. With respect to S138E, the intensity decreases during the first 24 h before stabilising until 72 h. Such tendency could be explained by a rearrangement of the two Trp residues in a more polar environment in the first steps of the aggregation process and the formation of turn intermediates observed on the CD spectrum. The fluorescence of Trp exhibits an increased sensitivity to collisional quenching, particularly due to adjacent groups within the protein, likely due to the propensity of the indole group in the excited state to transfer electrons [76]. While no shift in the emission band is observed in ITF spectra of S348E (Fig. 10C), its intensity decreases during the first 48 h, like S138E, before increasing within 72 h. This can be explained by conformational changes leading to Trp residues initially exposed to the solvent or polar sidechains in the first 48 h, followed by the formation of aggregates, within which the tryptophanyl moieties are buried in a hydrophobic core.

**Fig. 10.**
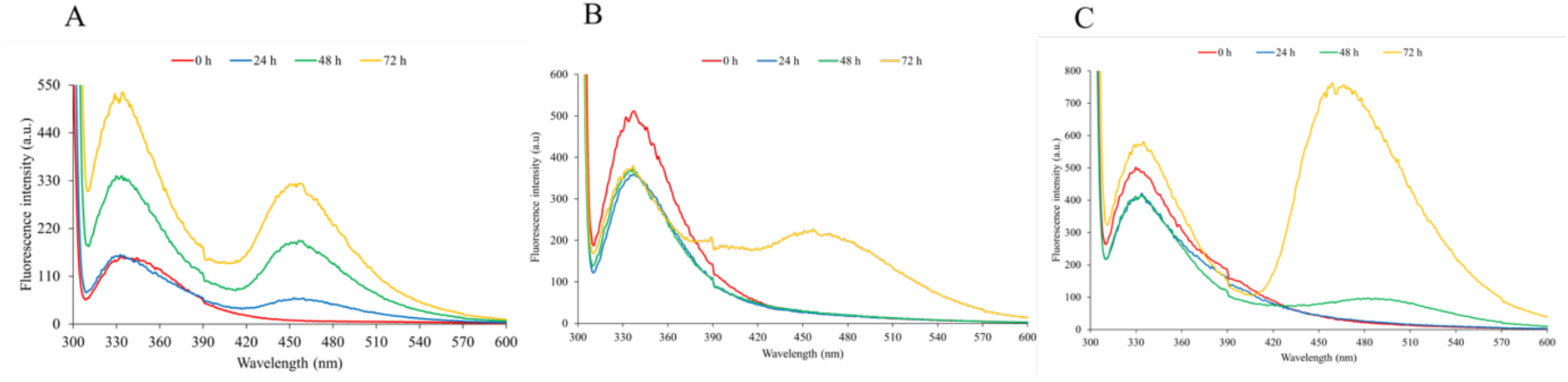
ITF spectra (λ_exc_ = 295 nm, sw = 10 nm) of (A) DPF3a WT, (B) S138E and (C) S348E after 0 h (red), 24 h (blue), 48 h (green), and 72 h (yellow) of incubation in TBS at ∼ 20 °C.

Such analysis is supported by the emergence of a second emission band at 456 nm, which has been assigned to the formation of amyloid fibrils for DPF3a and DPF3b [38,40,80]. Although its origins remain elusive, autofluorescence (AF) band could arise from low-energy electronic transitions induced by the delocalisation of electrons along hydrogen bonds between β-sheets [81–83]. AF could also originate from dipolar coupling between excited states of aromatic residues [84], change in amide groups communication and geometry [85], as well as charge transport or recombination [86]. While such AF band, designed as deep-blue autofluorescence (dbAF), is detected for WT DPF3a and the two phosphomimetics at the same wavelength, it emerges at different time scales and with varying intensities. Specifically, it appears after 72 h for S138E and 48 h for S348E, compared to 24 h for WT DPF3a, suggesting that both phosphomimetics aggregate at a slower rate than WT DPF3a. Furthermore, while its intensity is lower for S138E after 72 h compared to WT DPF3a, it is significantly higher for S348E, indicating that S138E likely forms fewer and/or smaller fibrils, whereas S348E leads to more and/or larger fibrils.

Concerning the ITyrF spectra of S138E (Fig. 11B) and S348E (Fig. 11C), they display similar trends regarding the time evolution of emission band intensity. The Trp-Tyr FRET emission band decreases in intensity in the first 24 h before increasing until 72 h of incubation, indicating that Tyr residues become less exposed to the polar environment, which is consistent with fibrillation and similar to WT DPF3a ITyrF spectra (Fig. 11A). Unlike S348E, which presents no spectral shift, but analogous to WT DPF3a, Trp-Tyr FRET emission band of S138E is blue-shifted from 337 to 335 nm, relative to less exposed residues. However, on the ITyrF spectra of S348E, the small shoulder at ∼305 nm, relative to Tyr residues, disappears over time, similar to WT DPF3a. Such behaviour indicates aggregation-drive conformational rearrangements of the protein, resulting in the enhancement of the FRET due to the closer proximity of Trp and Tyr residues. More significantly, the AF band at 456 nm is also present in the ITyrF spectra of all three proteins and is more intense than in ITF spectra, indicating not only that Tyr residues participate in AF but also contribute more to this intrinsic fluorescence. In line with the ITF spectra (Fig. 10), the dbAF band emerges after 48 h for S138E and 72 h for S348E, while it is detected as early as 24 h for WT DPF3a, confirming the phosphorylation-mediated kinetics discrepancy. After 72 h, a similar intensity pattern is observed, with S348E showing a higher intensity and S138E a lower one.

**Fig. 11.**
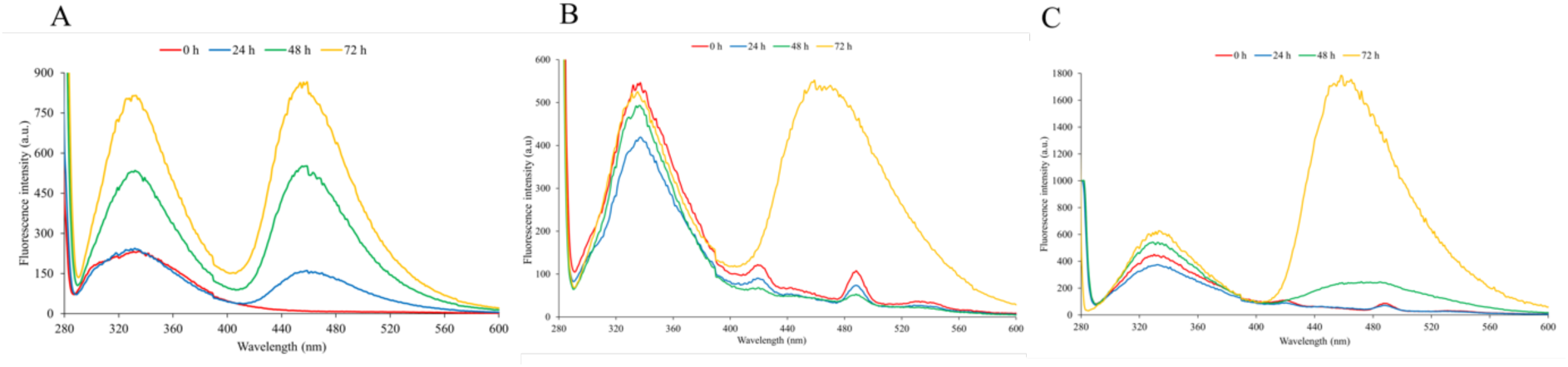
ITyrF spectra (λ_exc_ = 275 nm, sw = 10 nm) of (A) DPF3a WT, (B) S138E and (C) S348E after 0 h (red), 24 h (blue), 48 h (green), and 72 h (yellow) of incubation in TBS at ∼ 20 °C.

Despite the contribution of these aromatic residues and the peptide bonds, the predominant excitation wavelength contributing to this dbAF is 400 nm. The three proteins of interest have been excited at this wavelength and emission spectra have been obtained (Figure S2). The kinetics of aggregation of the two phosphomimetics can be compared to the one of WT DPF3a by plotting dbAF intensity at 456 nm against the incubation time (Fig. 12). This results in the emergence of three sigmoidal kinetic curves, characteristic of amyloidogenic proteins [87]. Such a kinetic profile has already been reported for WT DPF3a and DPF3b C-terminal parts and consists of three distinct phases [40]. First, the lag phase is associated with structural modifications of the monomers, leading to their assembly into oligomers. Afterwards, these enter then in the elongation or growth phase, characterised by a sharp rise in fluorescence intensity, where protofibrils are formed. Finally, the stationary phase is characterised by a stabilisation of the fluorescence intensity corresponding to the formation of mature amyloid fibrils [87,88].

**Fig. 12.**
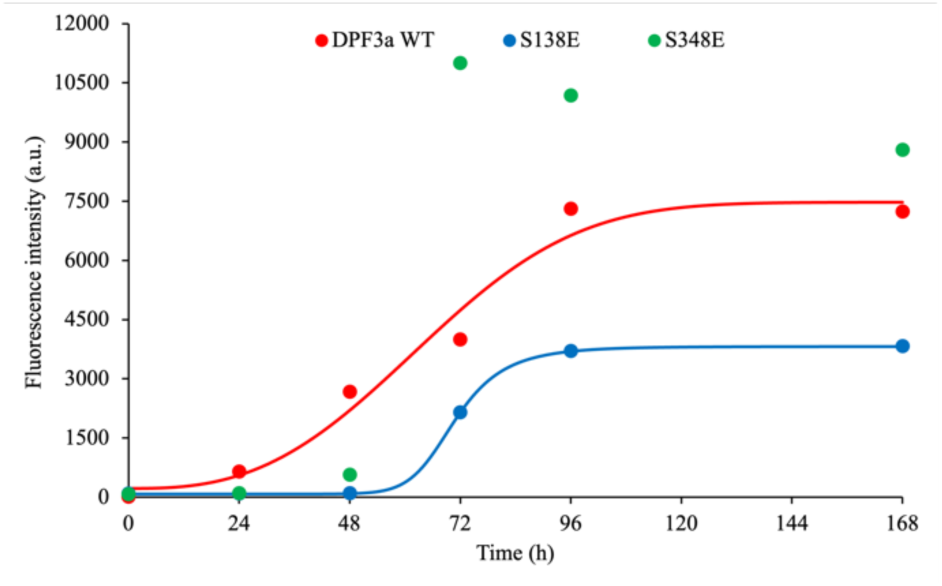
Sigmoid-fitted aggregation kinetic curves of DPF3b WT (red), S138E (blue) and S348E (green), using emission at 456 nm (λ_exc_ = 400 nm, sw = 10 nm). Samples were incubated in TBS ∼ 20 °C.

WT DPF3a presents the shortest lag phase, and the fluorescence intensity sharply increases until ∼96 h before reaching a plateau characteristic of the stationary phase. Regarding S138E, the lag phase is prolonged compared to WT DPF3a, which is in line with the slower increase in dbAF signatures observed for this phosphomimetic on ITF (Fig. 10B) and ITyrF (Fig. 11B) spectra. This is indicative of a delayed aggregation process. The elongation phase is comparatively shorter, giving way to the stationary phase with the plateau occurring at an intensity more than twice as low as that of WT DPF3a. For S348E, although a clear trend could not be established, the lag phase lasts up to ∼48 h, which is also longer than for WT DPF3a but faster than S138E, and is quickly followed by a sharp growth phase, with the intensity rapidly increasing within 24 h. The stationary phase therefore appears more rapidly than for the other two proteins. Like S138E, aggregation is delayed but the intensity of the plateau associated with the stationary phase is higher compared to WT DPF3a. This is in agreement with the high intensity observed on ITF (Fig. 10C) and ITyrF (Fig. 11C) spectra, and it likely reflects the formation of more fibrils and/or longer fibrils. The opposite is observed for S138E, which forms fewer and/or shorter amyloid filaments. The intensity of AF is low for S348E and almost negligible for S138E after 48 h of incubation, whereas CD spectra (Fig. 9B and 9C) indicate an enrichment in β-sheets structures. This can be explained by the formation of oligomers enriched in such structures that do not yet exhibit AF properties, emerging from a fibrillar organisation. These results demonstrate that the phosphomimetics, in addition to exhibiting a distinct aggregation mechanism compared to the WT, also follow different aggregation kinetics. This phenomenon has previously been observed for tau, where phosphorylation at multiple Tyr residues significantly delayed its aggregation [22]. The slower fibrillation kinetics for both phosphomimetics could be explained by the formation of the turn intermediates, as already observed in the case of amyloid β monomers, where the introduction of more constrained or stronger hydrogen-bonded β-turns significantly reduces or even inhibits fibrillation [89].

To further investigate the AF properties of the two phosphomimetics, excitation-emission matrices (EEM) were recorded after 7 (168 h) and 14 days of incubation. On the EEM of S138E (Fig. 13A) and S348E (Fig. 13C) recorded after 7 days, the dbAF band, which has already been visualised on ITF and ITyrF spectra, appears at an excitation and emission wavelength of ∼400 and ∼456 nm, respectively (∼400/456 nm). This originates from three contributions: the excitation of peptide bonds (λ_ex_ = 230 nm), the excitation of aromatic residues (λ_ex_ = 280 nm), and the primary contribution of dbAF (λ_ex_ = 400 nm). Interestingly, after 14 days, the intensity of the dbAF band of S138E (Fig. 13B) and S348E (Fig. 13D) decreases, while a second AF band emerges with a lower intensity. Specifically, this new population (∼350/400 nm) appears in the violet region, corresponding to violet AF (vAF). This AF mode has already been identified for DPF3a WT [80]. Once again, the intensity of the two AF populations (dbAF and vAF) is higher for S348E than for S138E, which is coherent with S348E forming a higher proportion of AF-emitting fibrillar species. The appearance of vAF after 14 days suggests structural rearrangements and the formation of aggregates with distinct structural properties compared to those observed after 7 days. Indeed, the interplay between AF populations may stem from the intrinsic dynamic nature of amyloid fibrils. The latter are known to exhibit polymorphism, meaning they can adopt multiple and distinct morphologies and stabilities despite being composed of the same protein [90]. These polymorphic aggregates undergo structural evolution over time, influenced by various parameters and governed by kinetics and thermodynamics principles [91]. The kinetically favoured amyloid fibrils are initially formed, followed by the transition to thermodynamically stable fibrils. The presence of polymorphism in amyloid fibrils is thought to play a role in the diverse biological effects associated with amyloid diseases [92]. Amyloid fibril polymorphism has already been reported for other amyloidogenic proteins like amyloid β, α-syn, and tau [93–95].

**Fig. 13.**
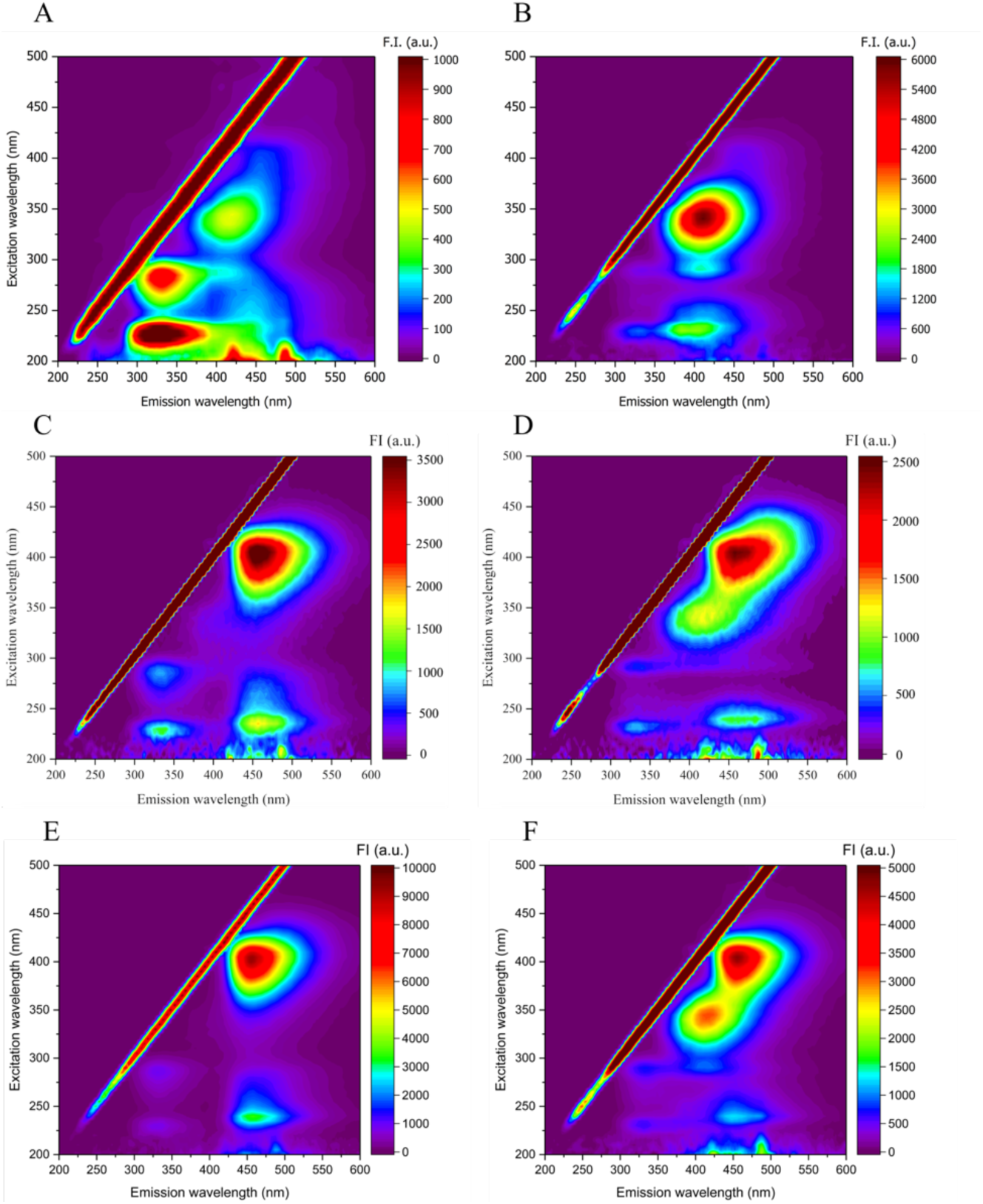
Excitation-emission matrices of S138E after (A) 7 and (B) 14 days and S348E after (C) 7 and (D) 14 days of incubation in TBS at ∼ 20 °C.

#### 3.3.2. Microscopic analysis of DPF3a phosphomimetics aggregate morphologies

By assessing the morphology of S138E and S348E aggregates by transmission electron microscopy (TEM) after 7 and 14 days, we could confirm that both phosphomimetics aggregate into amyloid fibrils with various morphologies. Regarding S138E after 7 days of incubation (Fig. 14A-B), 20 nm wide amyloid fibrils (black arrow) and amorphous aggregates (white arrow) are found entangled in fibrous networks (Fig. 14A). Straight fibrils (SFs) with a width ranging from 25 to 33 nm, which are larger than the 18 to 25 nm SFs formed by WT DPF3a [38], are also observed, reflecting the morphological variety of fibrils (Fig. 14B and Fig. S3A). Interestingly, after 14 days, double twisted fibrils (DTFs), with a width of 50 nm (black arrow) and a twist diameter ranging from 25 to 30 nm, are also found within amorphous phase (white arrow) (Fig. 14C and Fig. S3B). The formation of such twisted fibrils has previously been observed for WT DPF3a as early as 7 days of incubation, confirming a distinct and/or slower fibrillation process for S138E. Regarding S348E, while 30 nm wide SFs are also observed after 7 days of incubation (Fig. 14D), untwisted and curved fibrils (UCFs) of the same size appear intertwined, forming a fibrillar network (Fig. 14E and Fig. S3C). Both phosphomimetics appear to follow a fibrillation mechanism distinct from the one of WT protein, with each phosphomimetic exhibiting unique fibril morphologies, highlighting their structural diversity and polymorphic nature. Fibril polymorphism can result from the differences in the number and the arrangement of protofilaments composing the mature fibril, as well as the variation in the protofibril substructure [96]. Interestingly, after 14 days of incubation, S348E does not form DTFs as observed for S138E, but rather aggregates into accordion-like fibrils (ALFs) (Fig. 14F and Fig. S3D). This suggests a structural transition in the fibrils compared to those detected after 7 days, similar to what was observed for S138E, with a morphological diversity still evident after 14 days of incubation. The formation of structurally distinct aggregates over time could be relative to the emergence of the vAF population on the EEM (Figs. 13B-D). Given this morphological variety, it is not possible to correlate a single AF population with a unique fibrillar structure. However, for both phosphomimetics, the fibrils formed after 14 days, including the DTFs and ALFs, would represent the thermodynamically favourable fibrils, while the SFs and UCFs would correspond to the kinetically favourable ones.

**Fig. 14.**
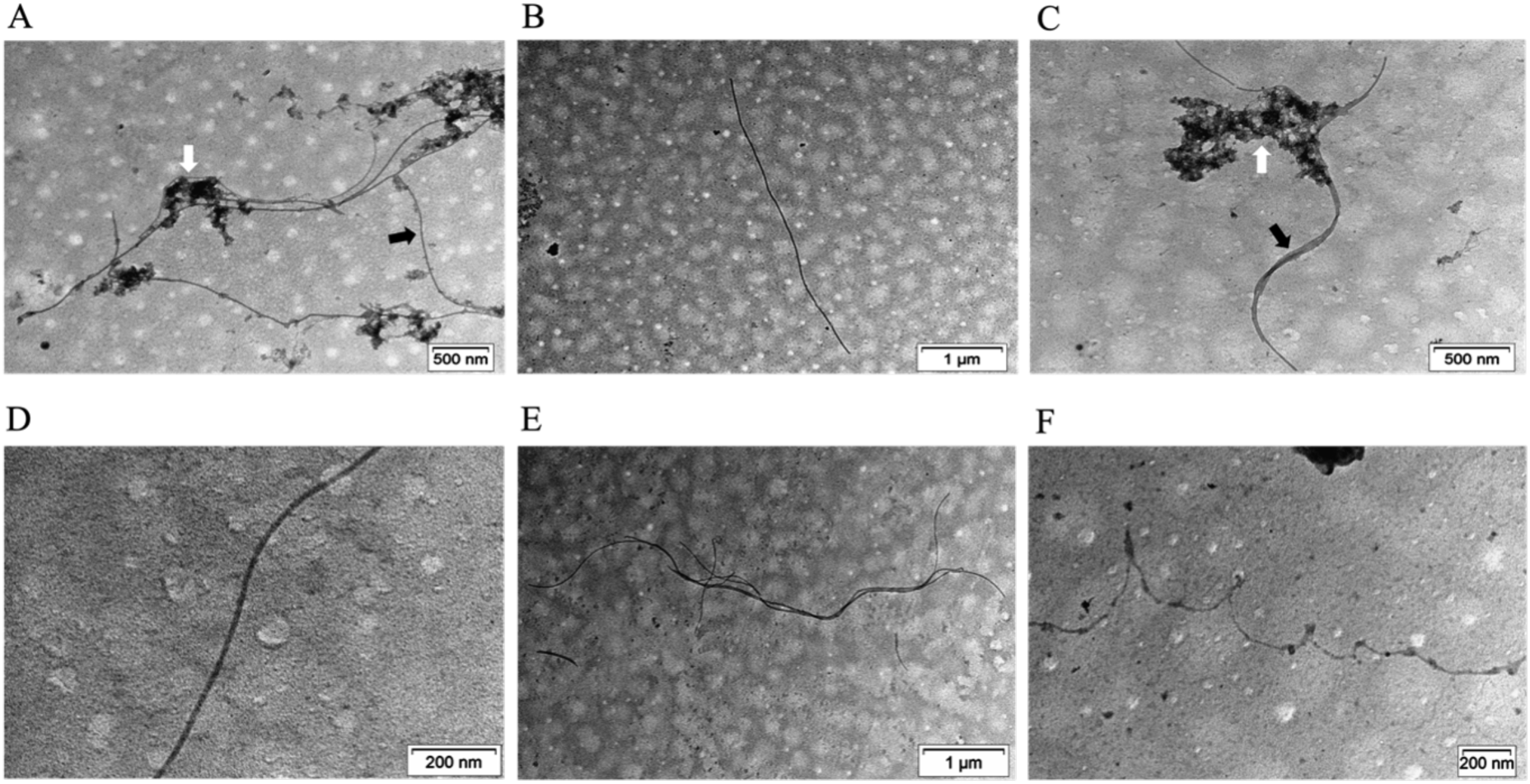
NS TEM micrographs (voltage of 100 kV) of S138E after (A-B) 7 and (C) 14 days and S348E after (D-E) 7 and (F) 14 days of incubation in TBS at ∼ 20 °C. (A) Amyloid fibrils (indicated by the black arrow) entangled in amorphous aggregates (indicated by white arrow). (B) Straight fibrils (SFs). (C) Double twisted fibrils (DTFs) (indicated by black arrow) and amorphous aggregates (indicated by white arrow). (D) SFs. (E) Untwisted and curved fibrils (UCFs). (F) Accordion-like fibrils (ALFs). The scale bar is placed at the bottom right of each micrograph.

The high resolution of atomic force microscopy coupled with infrared spectroscopy (AFM-IR) allows the visualisation of amyloid fibrils smaller than those observed by TEM [97,98]. AFM-IR is employed to gain further insights into the morphology and the secondary structure composition of WT DPF3a and both phosphomimetic aggregates after 14 days of incubation. While the three proteins assemble into elongated filaments upon aggregation, no significant morphological differences are observed amongst their smaller fibrillar assemblies. The latter, likely corresponding to protofibrils, appear structurally similar and may serve as the foundational units for the larger amyloid fibrils observed on TEM micrographs. The morphological differences thus likely arise at the level of these larger fibrils, while the initial protofibrillar structures remain conserved among the three proteins. However, the amyloid core is likely different for the two phosphomimetics compared to WT DPF3a, which could explain the AF discrepancies and the emergence of distinct fluorophores. Both WT DPF3a (Fig. 15A) and S138E (Fig. 15B) form fibrils with diameters of approximately 6-7 nm, accompanied by spherical oligomers of comparable or slightly smaller size. Additionally, larger fibrils, with heights reaching 11 nm for WT DPF3a (Fig. S4A) and 13 nm for S138E (Fig. S4B), have been observed. S348E, however, aggregates into protofibrils with a smaller size (approximately 4 nm), along with oligomers (Fig. 15C), as well as larger fibrils reaching up to 13 nm (Fig. S4C). Fibrils of this size have previously been observed for various isoforms of tau [99]. Finally, the strong IR signal at 1630 cm^-1^ (Fig. 15D-F and Fig. S4D-F) characteristic of β-sheet structures, is detected for each aggregate, confirming the fibrillation into β-sheet-rich amyloid structure.

**Fig. 15.**
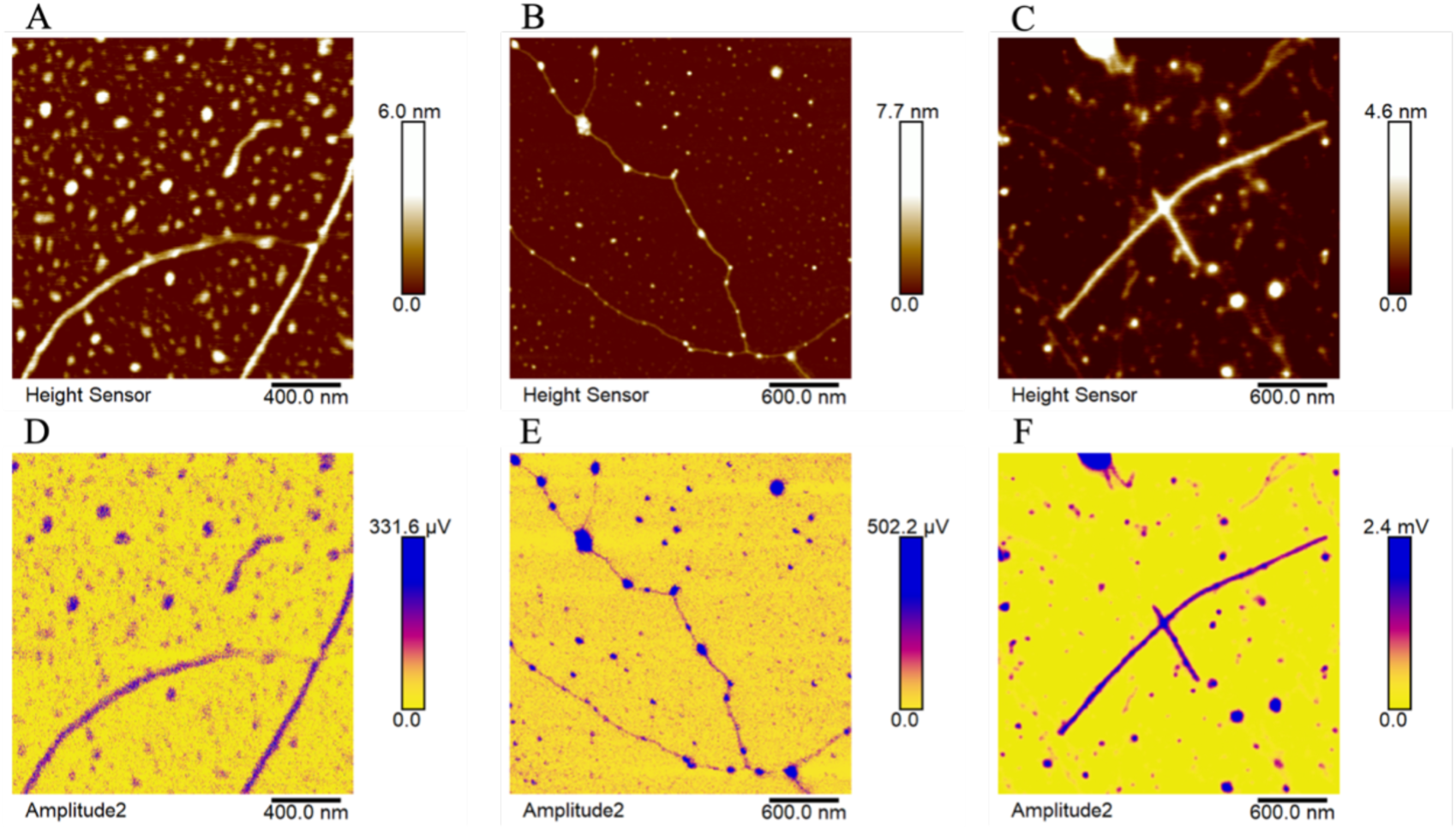
AFM micrographs of (A) DPF3a WT, (B) S138E, and (C) S348E after 14 days incubation in TBS at ∼ 20 °C. The scale bar is placed on the bottom right of each micrograph and the height on the right of each AFM micrograph. IR maps at 1630 cm-1 obtained from the AFM images of (D) DPF3a WT, (E) S138E, and (F) S348E.

In light of these results, in addition to exhibiting slower aggregation kinetics, the phosphomimetics display a distinct fibrillation mechanism compared to WT DPF3a, involving the formation of different intermediates and amyloid fibrils with distinct morphologies over time.

## 4. Conclusions

The present study has provided the first insights into the effect of phosphorylation on DPF3a structural and aggregation properties, by combining *in vitro* and *in silico* approaches. Far-UV CD spectroscopy has revealed typical spectral profiles of hybrid IDP for both phosphomimetics, with an increased turn and antiparallel β-sheet content, compared to WT DPF3a. ITF and ITyrF spectra have highlighted that Trp and Tyr residues in the phosphomimetics are partially solvent-exposed, as observed for WT DPF3a, although they appear slightly more buried, suggesting a conformational rearrangement mediated by the introduction of additional negative charges, notably a more folded N-terminal region.

MD simulations mimicking the *in vitro* conditions are in good agreement with these results, as pS138 and pS348 have displayed RMSD values typical of highly dynamic polypeptide chains. Phosphorylation of S138 and S348 has induced local variations in flexibility, especially the N-terminal 2/3 domain becoming more rigid. Notably, the C_2_H_2_ ZnF domain and the C-terminal extremity of pS138 have exhibited an increased flexibility. Overall, both phosphorylated proteins have shown increased Rg and SASA values compared to WT DPF3a. These differences can be attributed to the conformational changes induced by phosphorylation, particularly the elongation of the C-terminal extremity, likely driven by electrostatic repulsion between the phosphoserine and neighbouring negatively charged residues. Although phosphorylation of S138 and S348 promotes a closer proximity with arginine and lysine residues, leading to local collapsing, a higher number of H bonds was observed for WT DPF3a, supporting a more compact structure for the unphosphorylated form. Phosphorylation of S138 and S348 induce extended conformations of DPF3a, disrupting interdomain interactions, particularly between IDRs, and leading to a more isolated IDR-3, while pS348 retains partial contacts between the C_2_H_2_ ZnF and IDR-1.

DPF3a being an amyloidogenic IDP, the aggregation propensity of S138E and S348E were also investigated, using spectroscopic and microscopic techniques. While far-UV CD spectroscopy has unravelled an enrichment in antiparallel β-sheet over time for the three DPF3a constructs, ITF and ITyrF spectra have shown local structural rearrangements with Trp and Tyr found in more buried environment upon aggregation. Although dbAF-emitting species have been identified for both phosphomimetics over time which is indicative of amyloid fibrils formation, they fibrillation kinetics differ from that of WT DPF3a. Both phosphomimetics fibrillate more slowly, probably due to the formation of turn intermediates, even though S348E have shown a higher dbAF intensity, associated with the formation of more fibrils and/or longer filaments.

Based on our spectroscopic and microscopic analysis, a fibrillation pathway for the three proteins may be proposed (Fig. 16). Initially, WT DPF3a forms intermediates enriched in α-helix, while both phosphomimetics, already enriched in β-sheets and turns, reassemble to form turn-enriched intermediates, indicating a distinct aggregation pathway for the phosphomimetics. As the aggregation progress, all three proteins continue to undergo conformational rearrangements, leading to an enrichment in antiparallel β-sheet structures, forming the amyloid core. This leads to the formation of oligomers, which subsequently assemble into protofibrils that elongate and further assemble into SFs, which are larger for S138E and S348E. After 7 days of incubation, S348E also aggregates into UCFs, illustrating a certain morphology diversity. After 14 days, S138E forms DTFs, while S348E assembles into ALFs, highlighting the time-dependent differences in the fibrillation of these phosphomimetics, which is also associated with the emergence of vAF.

**Fig. 16.**
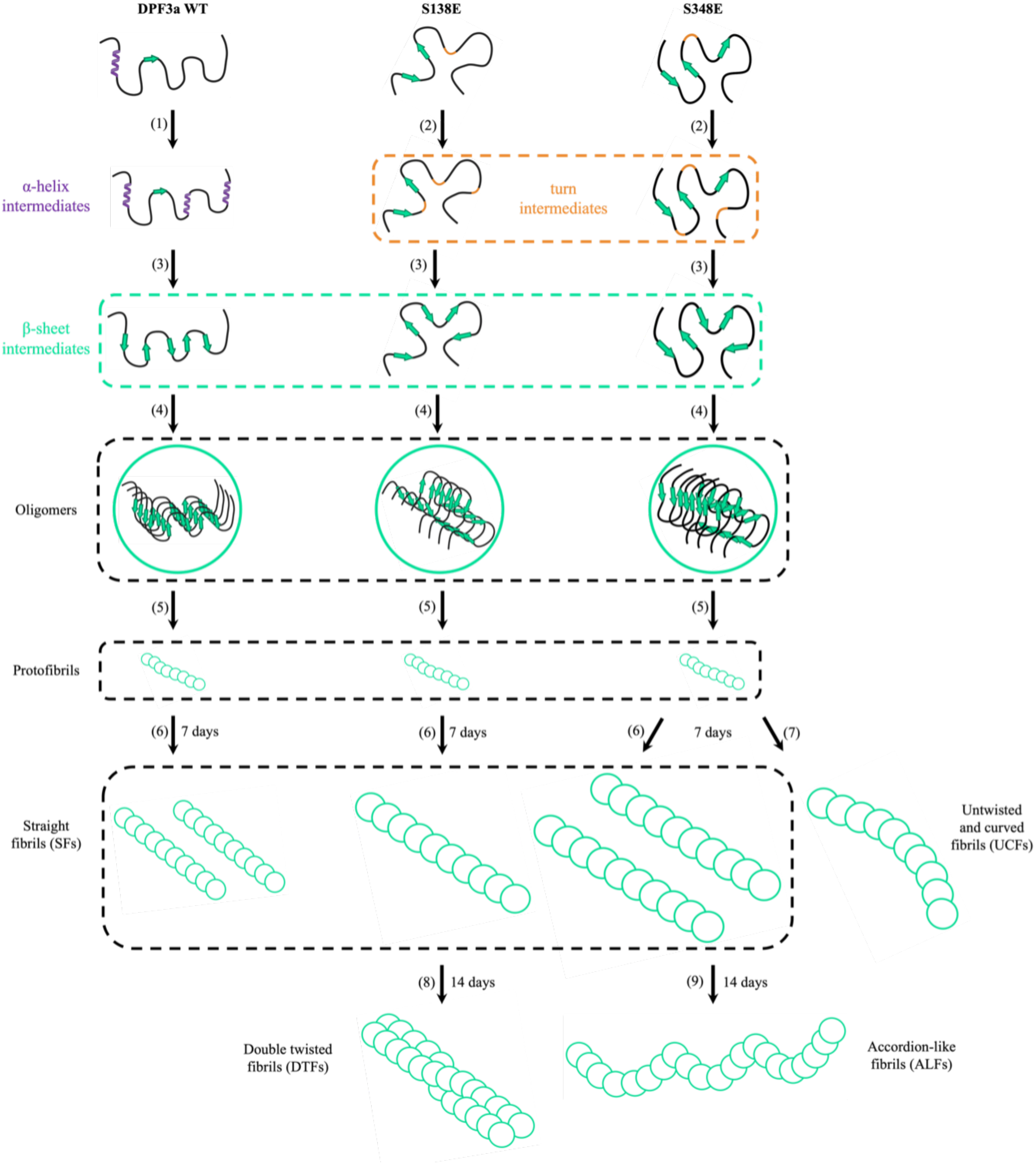
Suggested fibrillation pathways of DPF3a WT, S138E, and S348E. (1) DPF3a WT forms α-helix-enriched intermediates, while (2) both phosphomimetics are enriched in turn. (3) The three proteins continue to rearrange and become enriched in β-sheets, forming the amyloid core, (4) which leads to the formation of the monomers and (5) subsequently to the assembly of protofibrils. These elongate and assemble into SFs. (7) After 7 days, S348E also aggregates into UCFs. After 14 days, (8) S138E forms DTFs, while (9) S348E aggregates into ALFs.

Beyond the present study, further investigation would involve the *in vitro* phosphorylation of DPF3a by CK2, in order to assess the impact of multi-site phosphorylation on the structural conformation and aggregation propensity of the protein. Alternatively, a double phosphomimetic, such as the substitution of S138 and S348 with Glu residues, could be used to mimic the effects of CK2-mediated phosphorylation. A detailed understanding of the structural and aggregation features of phosphorylated DPF3a is indeed crucial for the structure-function of DPF3a and for the design of targeted anti-amyloid drugs.

## Abbreviations

α-syn: α-synuclein
AD: Alzheimer’s disease
AF: autofluorescence
AFM: atomic force microscopy
BAF: BRM/BRG1-associated factor
CD: circular dichroism
dbAF: deep-blue autofluorescence
d_ee_: end-to-end distance
DPF3: double plant homeodomain fingers 3
EEM: excitation-emission matrices
FRET: fluorescence resonance energy transfer
IDP: intrinsically disordered protein
IDR: intrinsically disordered region
IR: infrared spectroscopy
ITF: intrinsic tryptophan fluorescence
ITyrF: intrinsic tyrosine fluorescence
LCD: low complexity domain
MD: molecular dynamics
PD: Parkinson’s disease
PHD: plant homeodomain
PBS: phosphate-buffered saline
pDPF3a: phosphorylated DPF3a
PTM: posttranslational modification
R_g_: radius of gyration
RMSD: root-mean square deviation
RMSF: root-mean square fluctuation
SASA: solvent accessible surface area
TEM: transmission electron microscopy
TBS: Tris-buffered saline
vAF: violet autofluorescence
ZnF: zinc finger

## Data availability

Data will be made available from the corresponding author upon request.

## Acknowledgements and funding

The authors are grateful to the Research Unit in Biology of Microorganisms (URBM), as well as to the L.O.S. and Morph-Im platforms of the University of Namur. The authors are also appreciative of the PTCI high-performance computing resource of the University of Namur. The present research benefited from computational resources provided by the Consortium des Équipements de Calcul Intensif (CÉCI), funded by the Belgian National Fund for Scientific Research (F.R.S.-FNRS) under grant n°2.5020.11 and by the Walloon Region, and made available on Lucia, the Tier-1 supercomputer of the Walloon Region, infrastructure funded by the Walloon Region under the grant agreement n°1910247. T. L. and Q. M. thank the Fund for Research training in Industry and Agriculture (FRIA) Doctoral grant and Molecular-Scale Biophysics Research Infrastructure (MOSBRI), a European Union Horizon 2020 research and innovation program under the grant agreement N° 101004806. J. M. acknowledges the FNRS for its Research Fellow fellowship. The dimension IconIR used for AFM-IR experiment has been funded by the Walloon Recovery Plan (*Plan de Relance de la Wallonie* (PRW) – Belgium) in the framework of the Biogreen technological platform of excellence.

A. M. thank ANR and CGI for their financial support of this work through Labex SEAM ANR11 LABEX 086, ANR 11 IDEX 05 02. The support of the IdEx “Université Paris 2019” ANR-18-IDEX-0001 is also acknowledged. C. M. and D. M. also thank the FNRS for their Senior Research Associate position.

## CRediT authorship contributions

Tanguy Leyder: Conceptualisation; Methodology; Validation; Formal analysis; Investigation; Data acquisition and interpretation; Visualisation; Writing-original draft; Writing-reviewing & editing.

Julien Mignon: Methodology; Validation; Formal analysis; Data acquisition and interpretation; Writing-original draft; Writing-reviewing & editing.

Emma Bongiovanni: Validation; Data acquisition and interpretation; Writing-reviewing & editing.

Quentin Machiels: Validation; Data acquisition and interpretation; Writing-reviewing & editing.

Jehan Waeytens: Validation; Data acquisition and interpretation; Writing-reviewing & editing.

Vincent Raussens: Validation; Writing-reviewing & editing.

Antonio Monari: Conceptualisation; Validation; Data interpretation; Writing-reviewing & editing.

Denis Mottet: Validation; Writing-reviewing & editing.

Catherine Michaux: Conceptualisation; Validation; Supervision; Writing-reviewing & editing.

## Author information

Tanguy Leyder

Julien Mignon

Emma Bongiovanni

Quentin Machiels

Jehan Waeytens

Vincent Raussens

Antonio Monari

Denis Mottet

Catherine Michaux

