## supplementary information for "Unveiling the effect of phosphorylation on the structural and aggregation properties of the amyloidogenic intrinsically disordered protein DPF3a"

**–**

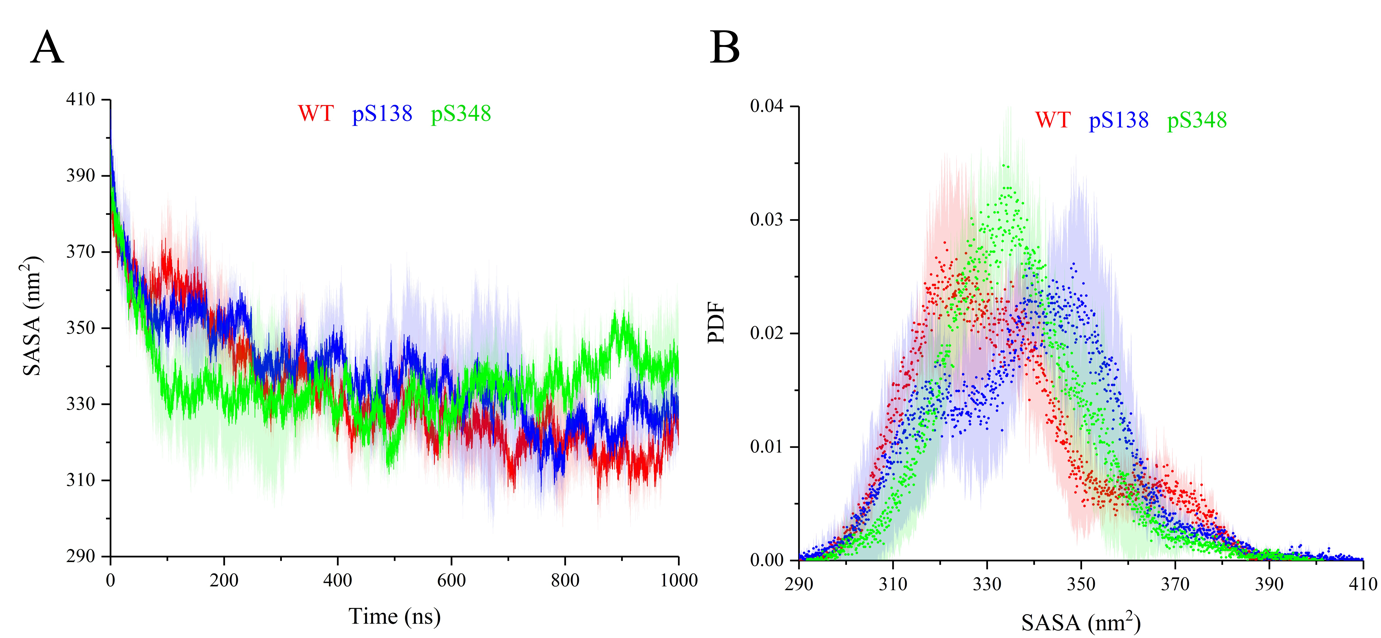

Figure S1 – (A) Time-evolution and (B) probability density function (PDF) distribution over the trajectories of the SASA of full-length WT (red), pS138 (blue), and pS348 (green) DPF3a simulated for 1 µs at pH 8.0 and 150 mM NaCl.

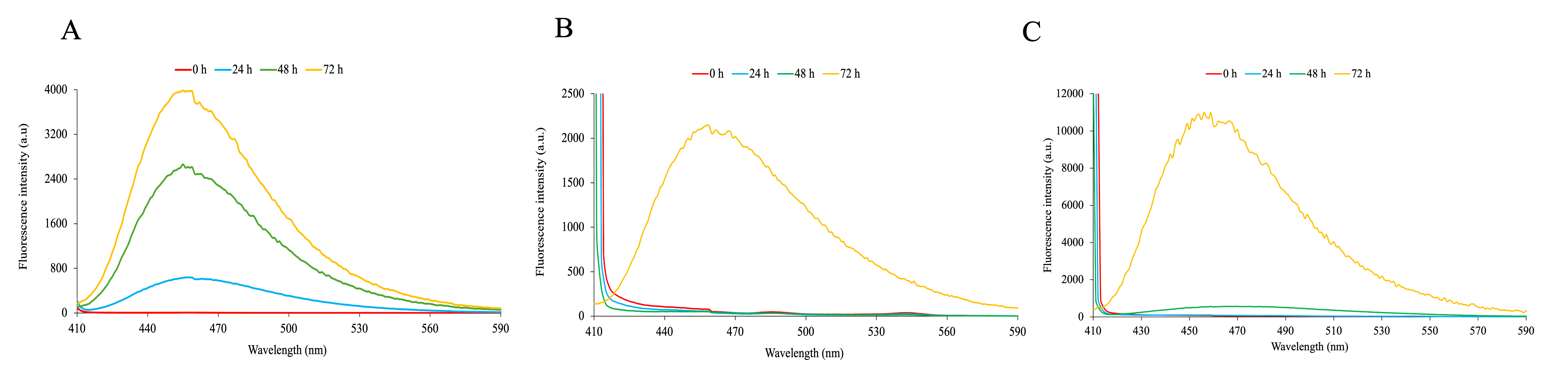

Figure S2 – Autoflluorescence spectra (λ_exc_ = 400 nm, sw = 10 nm) of (A) DPF3a WT, (B) S138E and (C) S348E after 0 h (red), 24 h (blue), 48 h (green), and 72 h (yellow) of incubation in TBS at ~ 20 °C.

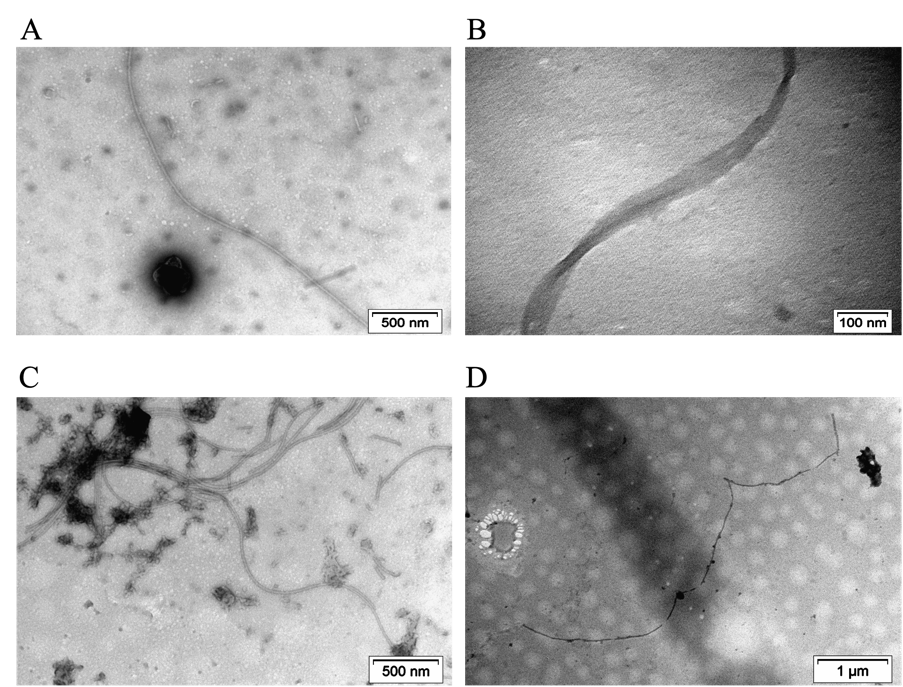

Figure S3 – NS TEM micrographs (voltage of 100 kV) of S138E after (A) 7 and (B) 14 days and S348E after (C) 7 and (E) 14 days of incubation in TBS at ~ 20 °C. (A) SFs. (B) Zoom on DTFs. (C) Network of UCFs. (D) ALFs.

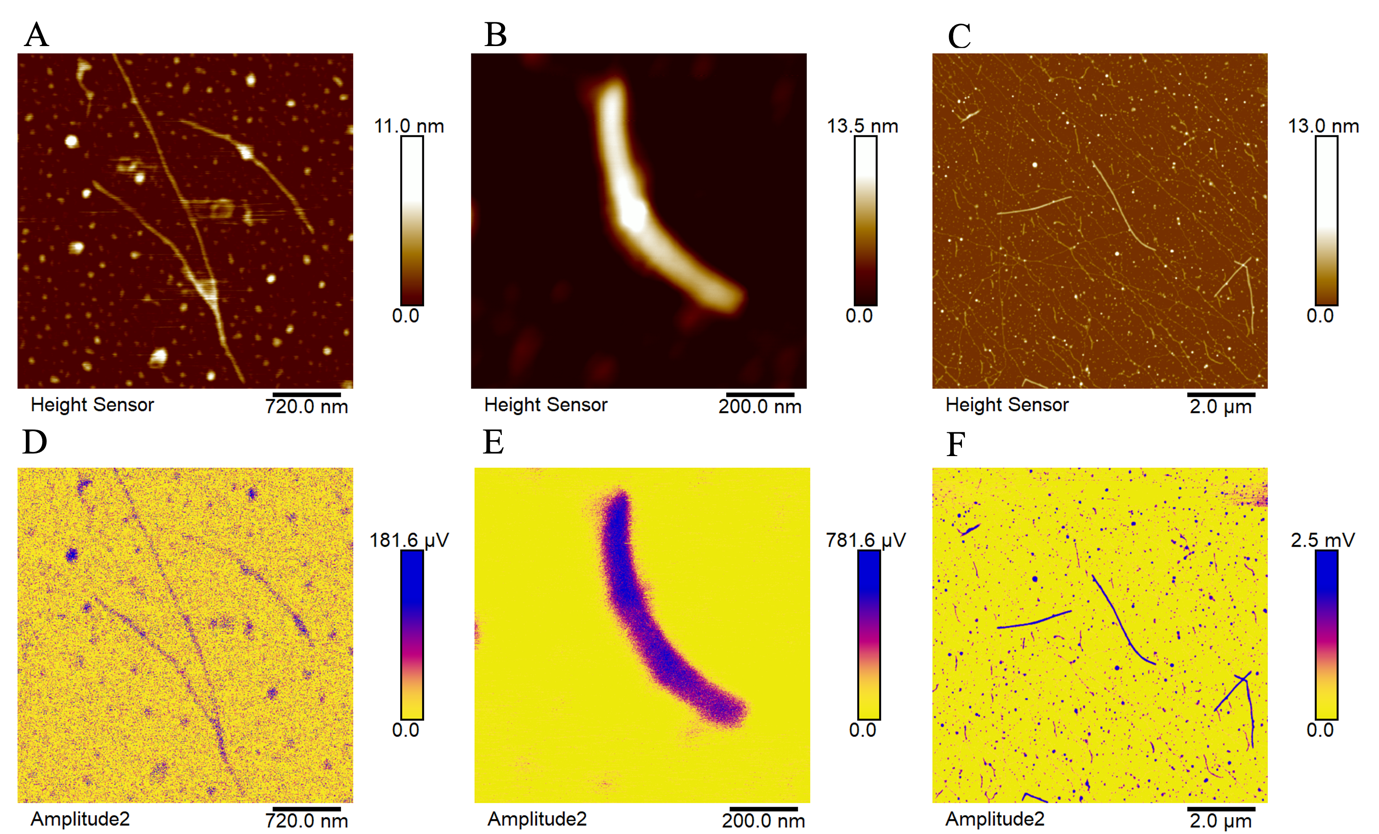

Figure S4 – AFM micrographs of (A) DPF3a WT, (B) S138E, and (C) S348E after 14 days incubation in TBS at ~ 20 °C. The scale bar is placed on the bottom right of each micrograph and the height on the right of each AFM micrograph. IR maps at 1630 cm^-1^ obtained from the AFM images of (D) DPF3a WT, (E) S138E, and (F) S348E.
